# PTFSpot: Deep co-learning on transcription factors and their binding regions attains impeccable universality in plants

**DOI:** 10.1101/2023.11.16.567355

**Authors:** Sagar Gupta, Veerbhan Kesarwani, Umesh Bhati, Jyoti, Ravi Shankar

**Affiliations:** Studio of Computational Biology & Bioinformatics, The Himalayan Centre for High-throughput Computational Biology, (HiCHiCoB, A BIC supported by DBT, India), Biotechnology Division, CSIR-Institute of Himalayan Bioresource Technology (CSIR-IHBT), Palampur (HP), 176061, India; Academy of Scientific and Innovative Research (AcSIR), Ghaziabad, Uttar Pradesh - 201002

**Keywords:** Transcription factor, Transcriptional regulation, Deep Learning, DenseNet, Protein Modelling, Transformers

## Abstract

Unlike animals, variability in transcription factors (TF) and their binding regions (TFBR) across the plants species is a major problem which most of the existing TFBR finding software fail to tackle, rendering them hardly of any use. This limitation has resulted into underdevelopment of plant regulatory research and rampant use of *Arabidopsis* like model species, generating misleading results. Here we report a revolutionary transformers based deep-learning approach, PTFSpot, which learns from TF structures and their binding regions co-variability to bring a universal TF-DNA interaction model to detect TFBR with complete freedom from TF and species specific models’ limitations. During a series of extensive benchmarking studies over multiple experimentally validated data, it not only outperformed the existing software by >30% lead, but also delivered consistently >90% accuracy even for those species and TF families which were never encountered during model building process. PTFSpot makes it possible now to accurately annotate TFBRs across any plant genome even in the total lack of any TF information, completely free from the bottlenecks of species and TF specific models.

## Introduction

Finding TF binding regions is central to understanding the transcriptional regulation across the genome. The rise of high-throughput technologies to detect such TF-DNA interactions like protein binding microarrays (PBM) [1], ChIP-Seq [2] and its various variants like ChIP-exo [3], and DAP-seq [4] has resulted into an explosion of DNA binding region data for various transcription factors [5]. To this date, there are approximately 1,28,467 ChIP-seq experiments reported at GEO/SRA for human alone. However, capturing all TF-DNA interactions through such experiments in any organism itself is a costly and impractical affair. One essentially requires some able computational approach to identify such TFBRs.

Unlike animals where human has been the main focus, plants have enormous number of species which define an extremely huge search space for possible experiments to detect TF-DNA interactions. If one compares the status of developments in plants with respect to animals, a huge gap is evident with hardly eight species of plants binding regions sequenced, covering merely ∼700 ChIP/DAP-seq experiments for selected TFs, mostly related to *Arabidopsis thaliana* and *Zea mays*. This lag in experimental data is equally reflected in terms of software resources and algorithms development for plant TFBR discovery. While for animal/human several software have been developed, there has been a very limited developments for plants. **Table 1** lists some software available for animals and plants where clear skew is visible (more information available in **Supplementary Table 1**). Therefore, it becomes urgent to develop computational approaches which could model TF-DNA interactions accurately for plants system, which may also reduce the dependence on costly binding experiments to a great extent.

**Table 1.**
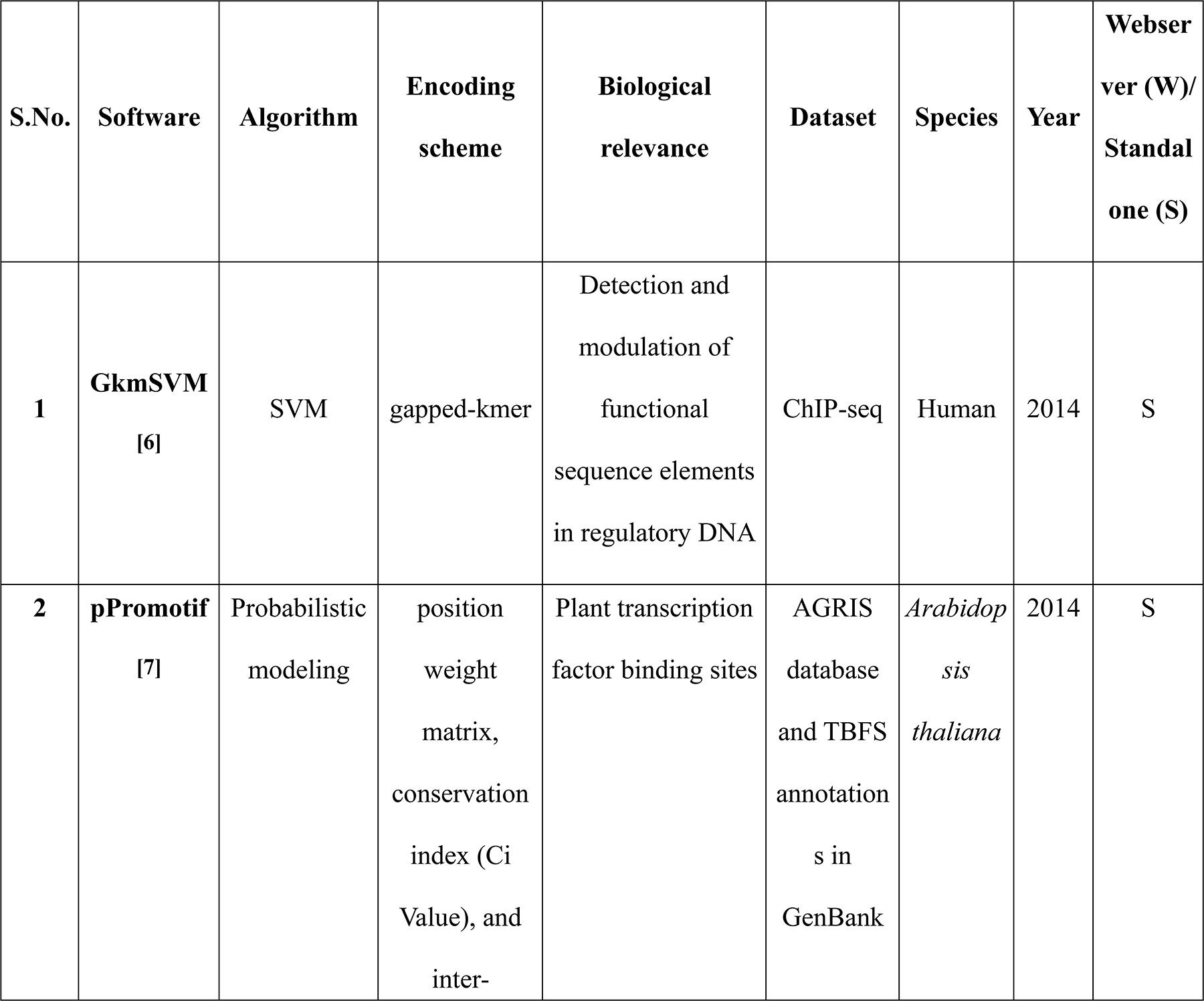
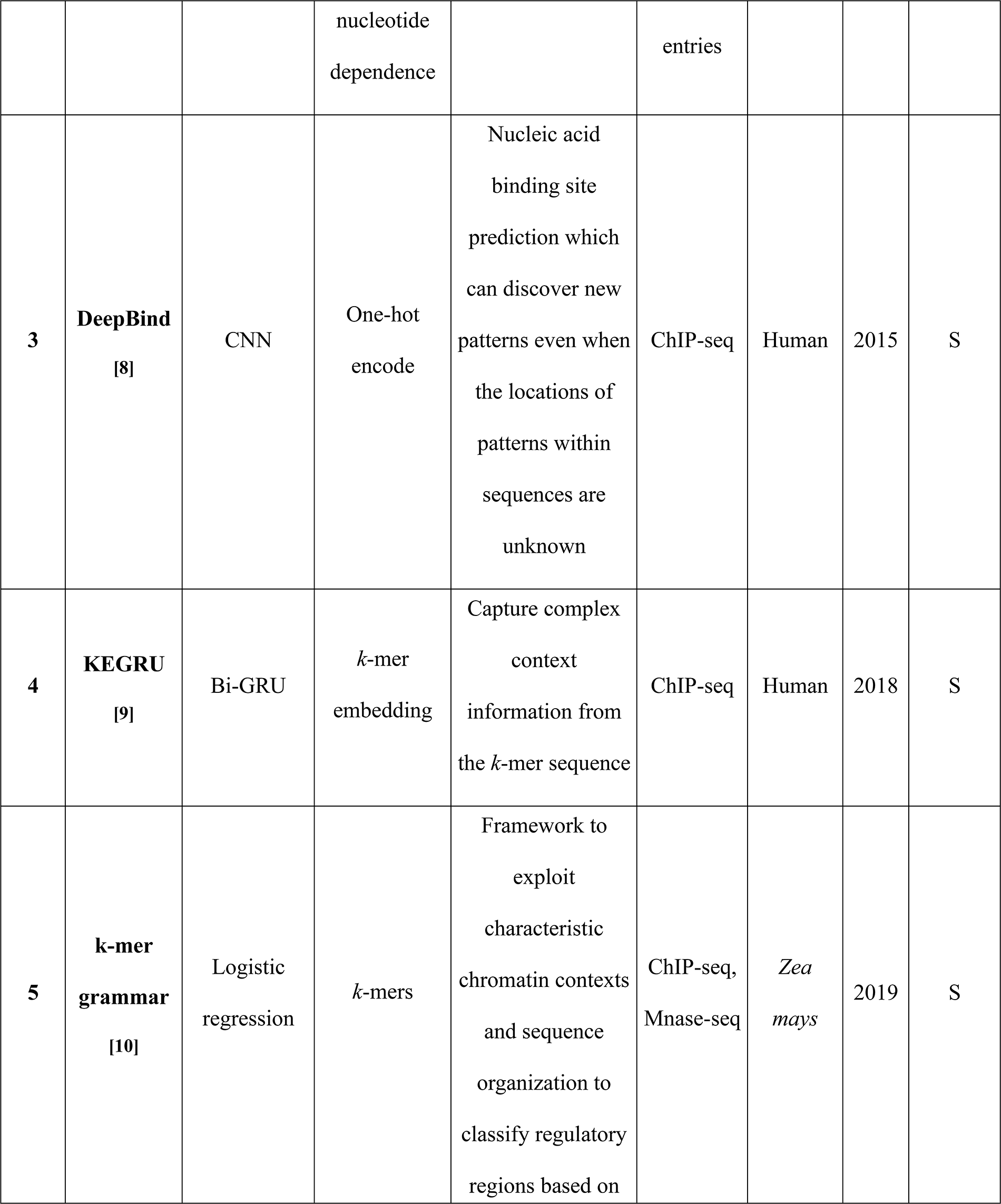
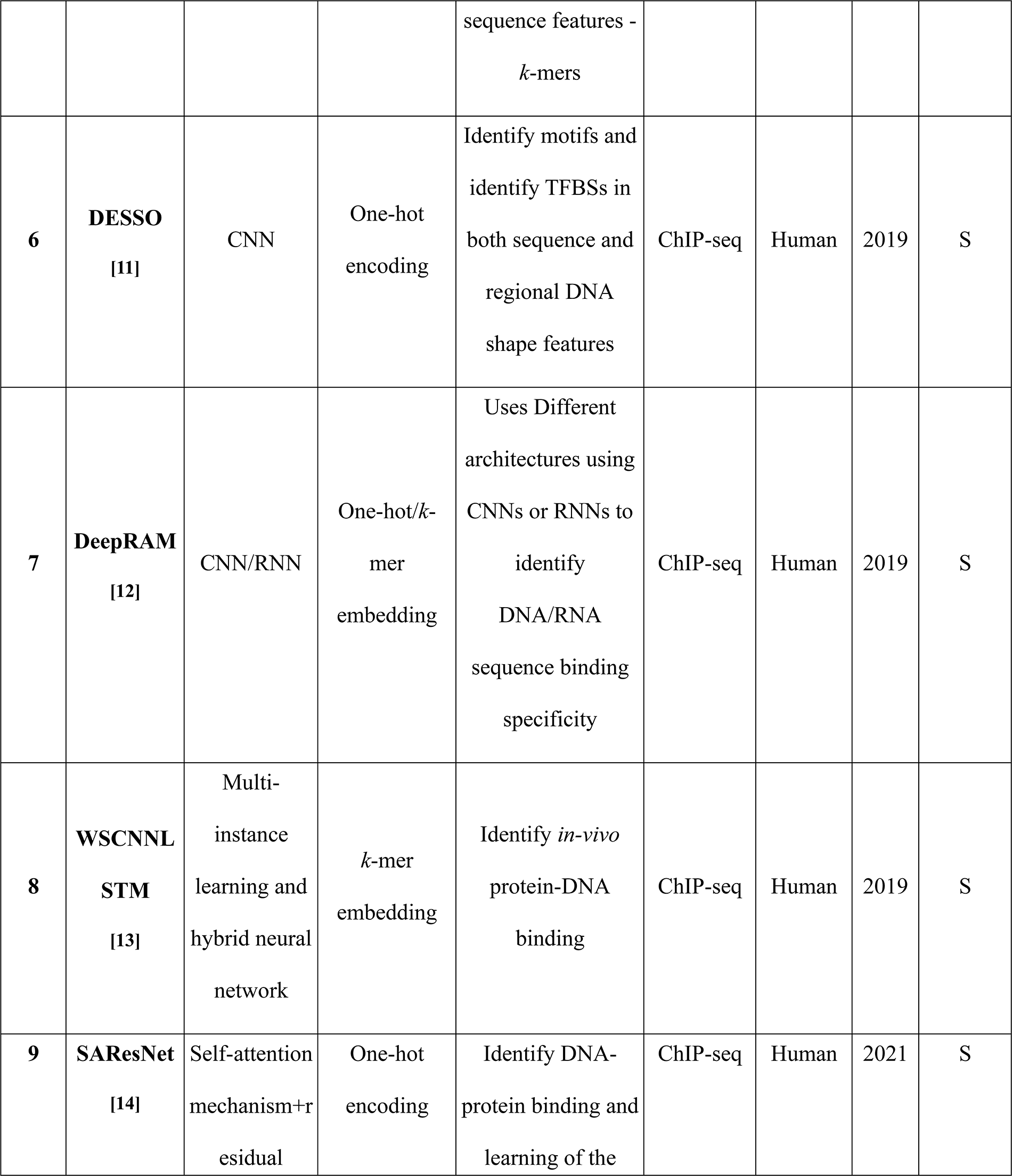
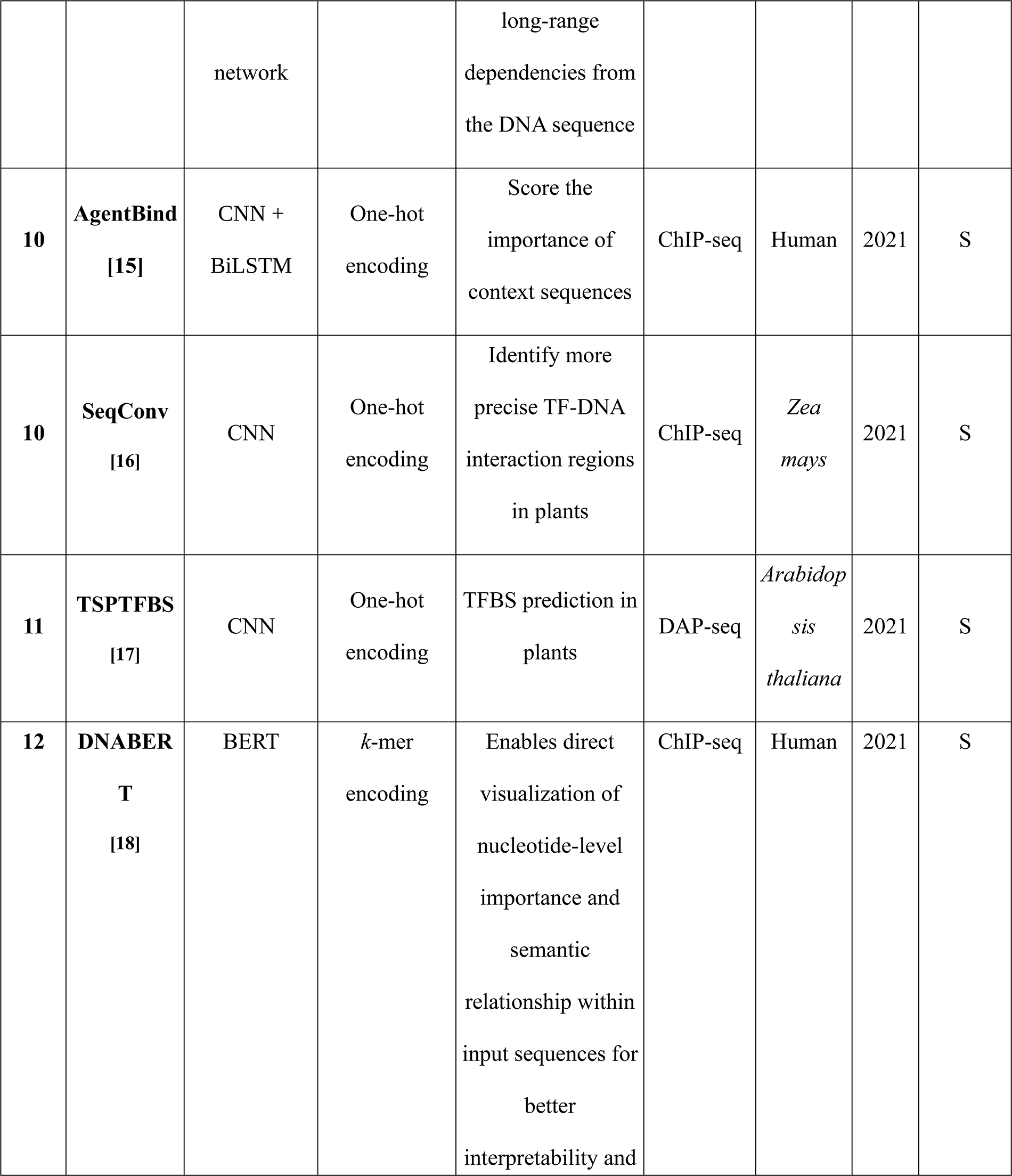
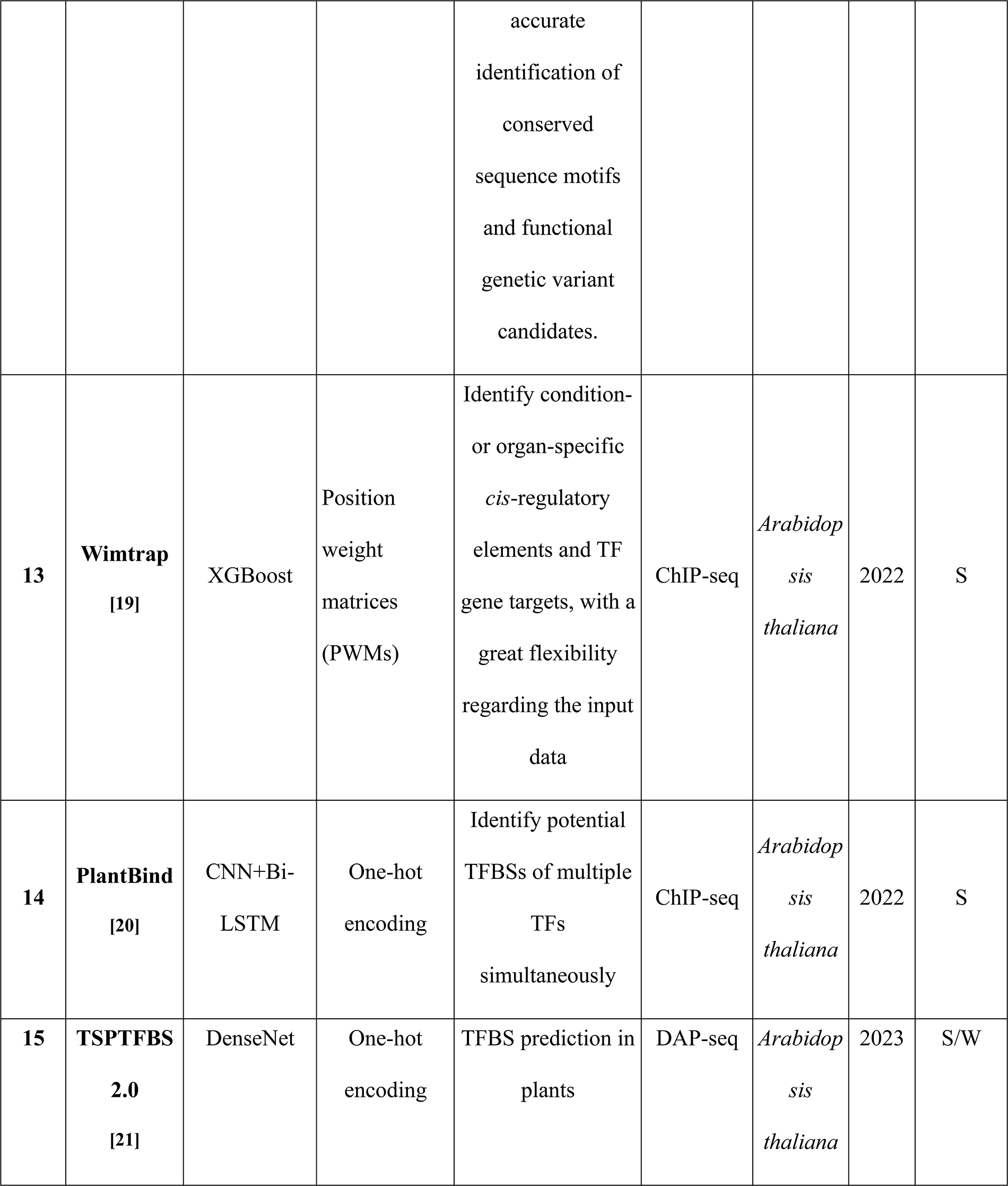
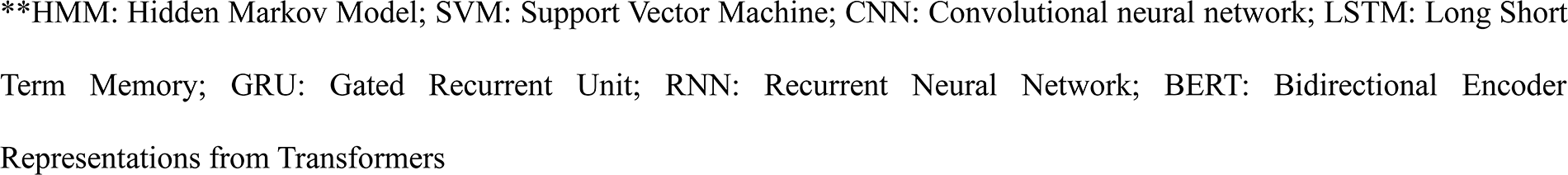
Brief list of some of the published tools for transcription factor binding site identification.

The existing software tools are overtly dependent upon the old school of motif discovery and user defined motifs, while reports suggest that TF binding is more about context and surroundings [17, 22–24]. The context and surroundings around an active binding motifs are defined by local sequence and shape preferences in highly specific manner. The motif finding step itself is heavily dependent upon binding experiments like PBM and DAP/ChIP-seq, on whose results, binding motif is defined for any given TF. Many TFs share a similar binding motif but yet differ in their binding due to local surroundings, shape, and contexts (**Figure 1a and b**) [10, 18, 20, 24]. Binding of a TF to its DNA target involves a prior step of local scanning of the target region in the range of around 90-150 bases. The sequence compositions, degenerate consensus sequences and cooperative motifs in the entire region were found contributing to the final halting of the TF around its target gene [25, 26]. Further to this, the choice of negative datasets with most of the software have been very relaxed as they randomly pick sequences and give too much weight to the consensus motif which can actually occur even in the non binding regions, creating weak datasets on which learning have been done so far. Thus, giving so much weight to some motif alone to determine binding of a TF to DNA is itself a misplaced practice.

**Figure 1:**
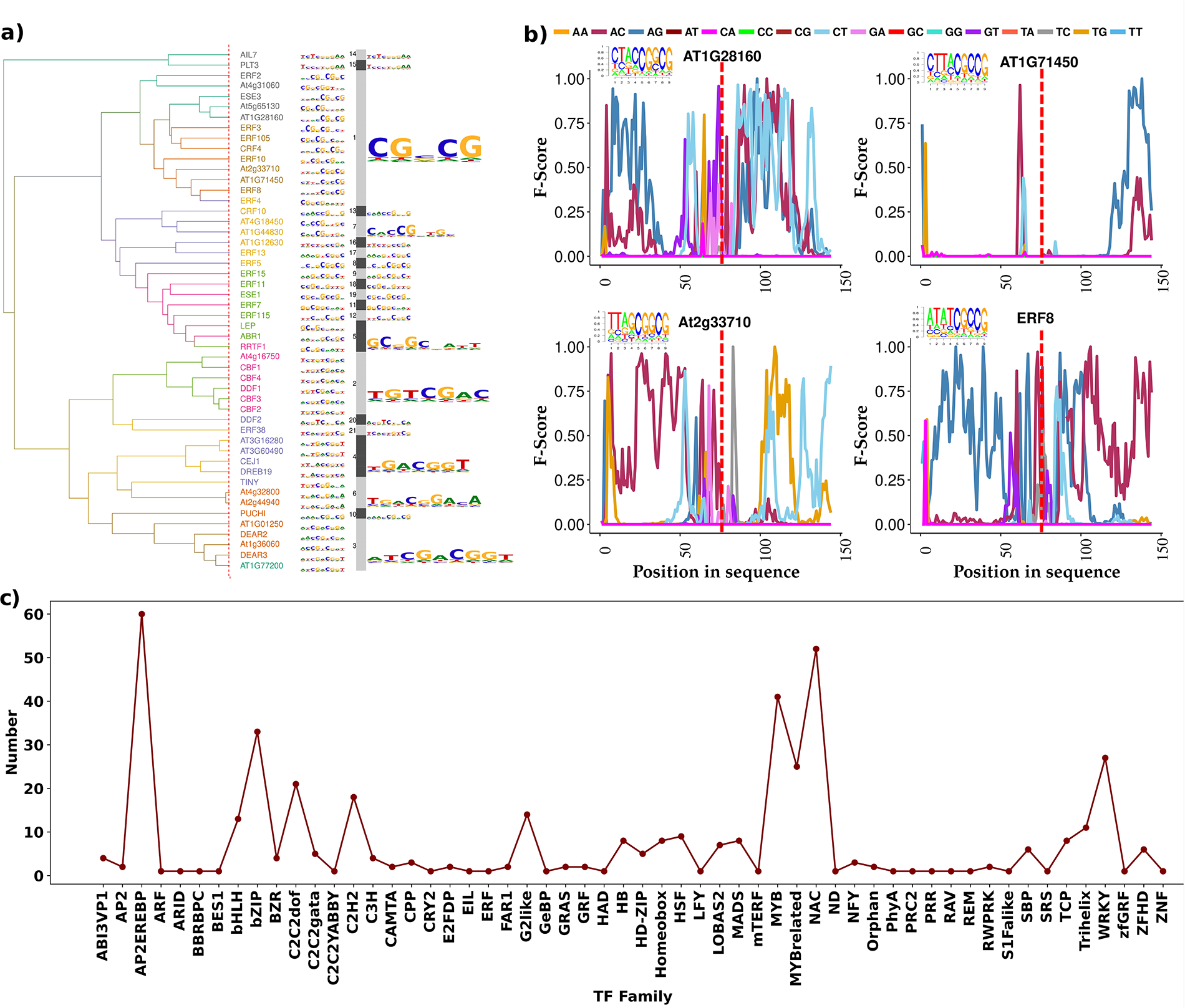
Motif and data. **(a)** Clustering plot of the prime motifs for TF family AP2EREBP. The TFs share similarity in their prime motifs. **(b)** Different characteristics spectra for TFs belonging to same family, same cluster, and sharing similar prime motif. This highlights the importance of context. Despite having similar binding motifs, their binding preferences differ from each other due to context. **(c)** Lineplot for the data abundance for the collected 441 TFs (ChIP and DAP-Seq) across 56 different family.

The most alarming matter is the fact that the plant biologists are overtly using TF and species-specific model like those developed for *Arabidopsis* specific TFs to report TF DNA binding in other species, attempting to answer transcriptional regulation misdirected, generating potentially misleading results and information. Its root is the lack of studies, data for other species, and lack of reliable computational resources specific to plant systems. Plant genomes in general exhibit enormous variability [27, 28], and transcription factors and their binding regions display large degree of variability across the plant species [29–32]. Thus, what may be working in *Arabidopsis* may not work in other plant species, and vice versa. Understanding and tackling the variability of plant transcription factors and their TFBRs is the main challenge to develop plant specific TFBR models where most of the currently existing software almost fail.

With this all foundations and understanding of the challenges, here present a unique and universal approach, PTFSpot, to detect TFBRs across plant genomes based on the following principles:

1) Instead of overtly relying upon motifs, identify the most significant motifs specific to any transcription factor and use it as the seed/anchor to identify most significant flanking regions for additional information. This is because the transcription factor scans a whole local region before halting at any given location, a process which depends a lot upon the flanking regions’ environment [25].

2) As already mentioned above, the enriched motifs are just one important feature. However, not all such motifs are bound by TF as the flanking regions’ environment is also important. And this information could become strongly discriminating if the negative set considers such unbound region’s motifs and flanking region information for them also. Therefore, use the significant motifs found in ChIP/DAP-seq data (positive datasets) to locate them in the unbound regions for the given experimental condition. This creates a highly confusing realistic negative dataset where overt importance of motifs is downplayed.

3) Using the motif seeds as anchors, represent the flanking regions with appropriate words of dimers (captures sequence composition as well as base stacking information), pentamers (reflects DNA shape), and heptamers (reflecting sequence motifs). Doing this may boost the discriminating power through sequence and structural information of the flanking regions.

4) Apply Transformers like state-of-the-art deep-learning algorithms which learn long distanced as well as local associations among the words, their co-occurrence while learning upon several hidden features, something not possible for traditional machine learning as well as other existing deep-learning methods.

5) Finding TFBR in cross-species manner fail because this is wrongly assumes that binding preferences of a TF remains static across the species while overtly relying on some enriched motif found in one or few species. While the fact is that the TF sequence, structure and its binding preferences vary across the species and even within their splice variants. Therefore, for reliable and universal TFBR discovery, one needs to learn the co-variability between the TF itself and its binding preferences. And if the available TFs and their binding preferences are learned this way together, they may bring a universal single model for TF:DNA interaction where co-variability between structures and sequences could answer binding preferences for any TF. Thus, learn co-variability between TF structure and binding regions while assuming all transcription factors under one hypothetical transcription factor which keeps changing its composition and structure (the corresponding transcription factor) according to which the preferred binding region also changes. This learning even from a single species may bring a universal model, which can work across any species even for never seen before transcription factors, as variability and interaction relationship is learned. Thus, learning this way even on *Arabidopsis* data alone, which is also the most abundant one, would turn to be a boon.

Based on these principles, we have developed a Transformer-DenseNet Deep-Learning universal system while learning from ChIP/DAP-Seq binding data for 436 TFs, and their corresponding 3D structural details. A highly extensive benchmarking study was carried out with three different experimentally validated datasets as well as never seen before species specific experimental binding datasets to test the universality of PTFSpot. The results have been ground-breaking with performance lead of >30% over the existing software pool. Also, in terms of cross-species performance, PTFSpot has delivered an outstanding performance where it consistently scored above 90% accuracy for never encountered before plant species and transcription factors. This is something which has never been witnessed before and revolutionary as it will empower to detect the TFBR across any plant species for any TF with impeccable accuracy and may even bypass the need of costly experiments like DAP-seq to detect the TFBRs.

## Materials and Methods

### Data retrieval

ChIP/DAP-seq peak data for 436 TFs spanning 5,753,198 distinct peaks were retrieved from PlantPAN3.0 (54 TFs) and Plant Cistrome Database (387 TFs) [33, 34]. Genomic sequences were extracted from the peak coordinates **(Supplementary Table 2 Sheet 1)**. All details on data retrieval are visualized in **Figure 2**.

**Figure 2:**
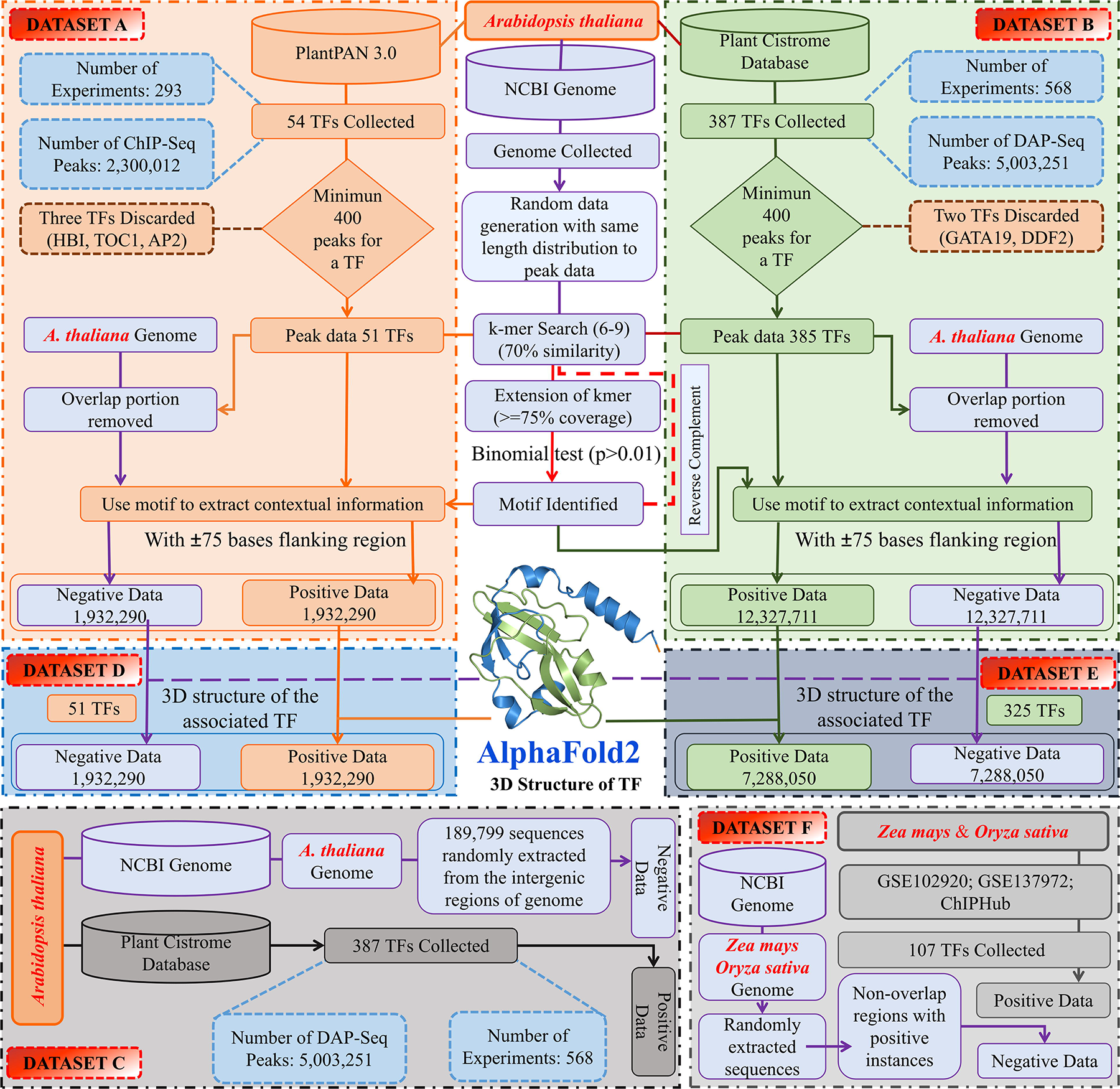
Flowchart representation of dataset formation. (**a)** The protocol followed for Dataset “A” creation, (**b)** Dataset “B” creation, **c)** Dataset “C” creation, (**d)** Dataset “D” creation, **e)** Dataset “E” creation, and (**f)** Dataset “F” creation. The datasets “A” and “B” contained the positive instances originating from ChIP-seq and DAP-seq, respectively, with sources PlantPAN3.0 and Plant Cistrome databases, respectively. Negative part of the datasets was formed by considering those genomic regions which display the regions similar to the prime motifs found enriched in DAP/ChIP-seq TF binding data but never appeared in the DAP/ChIP-seq data. Dataset “C” also contained the positive instances from the Plant Cistrome Database but for negative instances it contains random genomic regions. Most of the existing tools have used this dataset and its subsets. The datasets “D” and “E” were created from the datasets “A” and “B”, respectively, while adding up the TF structural data also. Dataset “F” was constructed to be used completely as a test set to evaluate the universal model and its applicability in cross-species manner. This dataset covered 117 TFs from *Zea mays* (93 TF) and *Oryza sativa* (24 TF).

### Approach to identifying motif seeds candidates and anchoring significant seeds

For the 436 *Arabidopsis thaliana* transcription factors, ChIP/DAP-seq peak regions were scanned to identify prevalent 6-mer seed candidates with ≥70% identity, based on previous observations [33–36]. Every sequence was initially represented in the form for overlapping hexameric seeds, which were used to scan across the sequences for the most similar seed regions among themselves and accordingly were piled up against each other. Enrichment of *k-mers* was determined by computing their occurrence probabilities in peak data relative to a random genomic background model following the peak length distribution. The seeds represented in ≥75% of peaks with significant enrichment (binomial test, p < 0.01) were selected and iteratively extended bidirectionally while retaining instances with ≥70% identity. For every extension step, statistical significance, the identity, and coverage criteria were repeatedly evaluated. The final motifs obtained after these steps were called the anchoring seeds which satisfied the two criteria: ≥75% abundance (p < 0.01) in the peak data and ≥70% identity. Most enriched ones among them were called the primes. They were used as the anchors to derive the flanking region contexts for the datasets creation. The process was done for both the strands. This motif discovery approach has been introduced by us previously [37] while the further details are given in the **Supplementary Information Materials and Methods section.**

### Dataset Creation

To generate the positive datasets, the peak data sequences were transformed into instances once the motifs were anchored for each TF in the provided peaks data. ±75 bases in both directions from the ends of the prime motif regions were taken to produce instances.

To form the negative datasets, regions reflected in the peak data were removed, leaving only unrepresented regions for selection. These regions were scanned for prime and reverse complementary motifs similar to the positive instances, flanked by 75 bases for context [38]. Positive and negative instances were pooled at a 1:1 ratio, without overlap.

Datasets “A” and “B” comprised positive instances derived from ChIP-seq and DAP-seq experiments, respectively, on *Arabidopsis thaliana* TFs. Dataset “C” leveraged DAP-seq data for 387 TFs from the Plant Cistrome Database [17, 21]. To develop universal models capturing TF-DNA interactions, Datasets “D” and “E” further incorporated the 3D structures of the corresponding TFs from AlphaFold2 [39]. Dataset “E”, with 325 *Arabidopsis* TFs, was used for training and evaluation. An independent test Dataset “F” was curated to assess cross-species generalization, comprising 117 TFs from *Zea mays* (93) and *Oryza sativ*a (24) with over 2.2 million ChIP-seq peaks from public data repositories [40]. For all the datasets every possible overlap and redundancies (full or partial) were screened out. Further details on dataset construction are provided in the **Supplementary Information Materials** and **Methods section** and **Figure 2**.

### Word representations and tokenization for sequence data

Utilizing a four-base DNA alphabet yielded 16 unique 2-mer words, 1,024 unique 5-mer words, and 16,384 unique 7-mer words. Dinucleotides, pentamers, and heptamers encode composition, base stacking, shape, and regional motif information [22, 37, 41]. The resulting word representations were tokenized by assigning a distinct integer to each unique word creating an input vectors of 469 tokenized words. These numeric token embeddings served as the inputs to the Transformer encoder part. The TensorFlow (Keras) tokenizer class was employed to implement the tokenization procedure.

### Implementation of the Transformers

Each tokenized sequence was converted into a two-dimensional word embedding matrix, with rows determined by the encoding vector size (*d*=28) and columns by the number of tokenized words (*n*). Positional encodings of dimension “*d”* were computed in parallel using sinusoidal functions [42] and combined with the word embeddings, forming the input M’ to the Transformer encoder. Within the multi-headed self-attention mechanism, the embedded sequence M’ was projected onto query (Q), key (K), and value (V) matrices using learnable weight matrices *W^Q^*, *W ^K^*, *W^V^*:

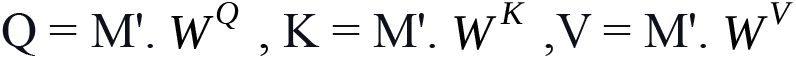

The attention scores were then computed through scaled dot-product attention between Q and K, followed by softmax normalization and multiplication with V:

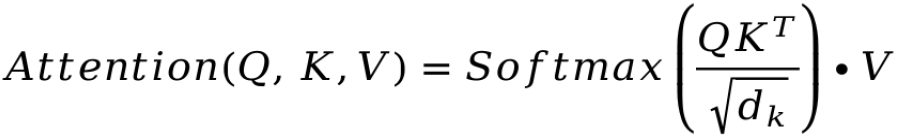

This computes pairwise attention weights between all words, capturing their contextual associations.

A multi-headed transformer featuring 14 attention heads was executed this process in parallel, with their outputs concatenated to capture different relational perspectives. The combined contextual representations underwent feed-forward processing, normalization, dropout regularization and global average pooling and a final classification layer with a sigmoid activation function for binary prediction of TF binding region. The Adam optimizer [43] was employed for model training using binary cross-entropy loss. This multi-headed self-attention architecture enabled effective learning of long-range dependencies and contextual TF-DNA binding preferences from the tokenized, embedded DNA sequence inputs across 436 *Arabidopsis thaliana* TFs. Hyperparameter optimization was conducted employing Bayesian optimization. Comprehensive implementation and optimization details of the transformer system are given in the **Supplementary Information Materials and Methods section**.

### Structural and molecular dynamics studies

The 3D structures of TFs were modeled using AlphaFold2 [39], with the top-ranked model selected for the study. ScanProsite (http://prosite.expas.org) confirmed the functional domain and amino acid residues in the active site pocket. Comparative studies were conducted based on protein sequence, functional domain, 3D structure, and binding affinity. Additional details regarding the validation of prime motif and transcription factor interactions through molecular docking [44] and simulation is given in the **Supplementary Materials and Methods section**.

### Transformer-DenseNet system for cross-species identification of the binding regions

The TF and binding regions covariability was learned through a hybrid Transformer-DenseNet system. DenseNet is a very high depth CNN based Deep-Learning architecture which learns the spatial patterns much efficiently than CNNs due to its higher depth and capability to keep the learning from previous layers afresh while effectively mitigating the vanishing gradient issue. 3D structure information, corresponding binding region from ChIP/DAP-seq data for each TF were considered for its training. The DenseNet [45] part processes atom-wise coordinates while the Transformer part processes the sequence-based input as described in the section above on Transformers. Normalized coordinates are inputted into a convolution layer with the dimensions of 300×24×3, accommodating the TF’s amino acid positions and atoms. Zero-padding maintains consistent matrix dimensions for shorter sequences. This approach stems from an analysis of over 400 TFs, revealing a maximum of 24 atoms per amino acid. Full implementation and optimization details are provided in the **Supplementary Materials and Methods section**.

### Building the DenseNet Architecture

Each layer in DenseNet receives information from all preceding layers, facilitating more efficient feature learning. DenseNet model employed in our study consists of one convolution layer with 32 filters (kernel size = 3), followed by batch normalization and 2D maxpooling (stride = 2). It comprises ten dense blocks and nine transition layers, totaling 121 layers.

Within each dense block, the input is concatenated with the feature maps of all preceding layers, denoted as (*m*_0_, *m*_1_, … *m_l_*_-1_,). Each layer within the dense block consists of batch normalization, ReLU activation, and 3x3 convolution. Transition layers facilitate down-sampling and include batch normalization, 3x3 convolution, maxpooling2D, batch normalization, another 3x3 convolution, and a dropout layer. The growth rate ’*k*’ determines the number of feature maps contributed to the global state.

After down-sampling, the output is flattened and concatenated with the transformer output for classification. This concatenated output undergoes batch normalization, dropout, dense, and dropout layers before passing through a sigmoid activation-based single-node classification layer. We utilized the “Adam” optimizer for weight adjustment, with a batch size of 64 and six epochs. Further details of this module is provided in the **Figure 3**.

**Figure 3:**
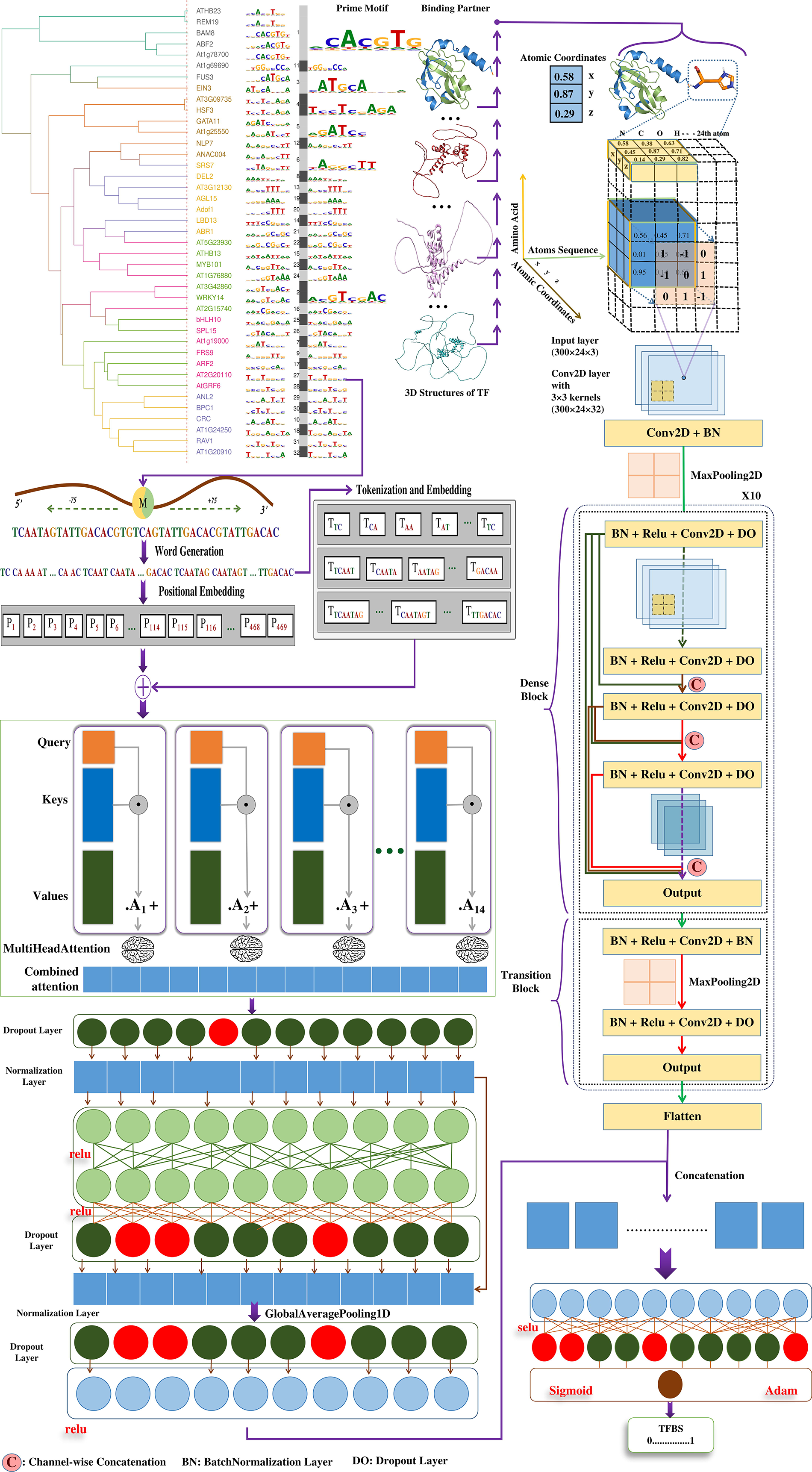
Implementation of the PTFSpot Deep Co-learning system using Transformers and DenseNet to identify TFBR across plant genomes. The first part is a 14 heads attention transformers which learn from the dimeric, pentameric and heptameric words representations of any given sequence arising from anchoring prime motif’s context. In the parallel, the bound TF’s structure is learned by the DenseNet. Learning by both partners are joined finally together, which is passed on to the final fully connected layers to generate the probability score for existence of binding region in the center.

### Performance Assessment

According to standard practice, every dataset involved in training-testing was divided into train (70%) and test datasets (30%). The developed Transformer-DenseNet model was tested on the 30% intact and completely untouched test portion. Four categories of performance confusion matrix namely true positives (TP), false negatives (FN), false positives (FP) and true negatives (TN) were evaluated. The performance of the built Transformer-DenseNet model was evaluated using performance metrics such as sensitivity, specificity, accuracy, F1 score, and Mathews correlation coefficient (MCC) [46].

Performance measures were done using the following equations:

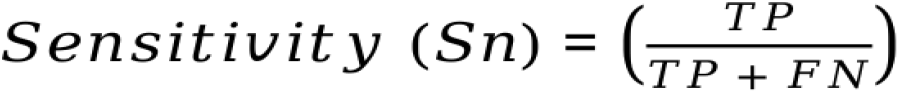

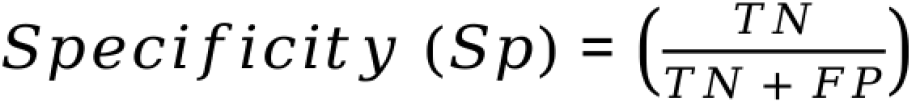

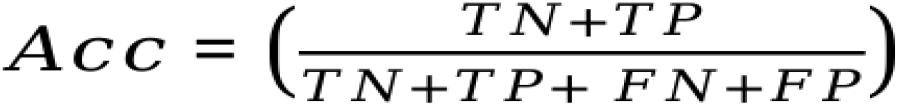

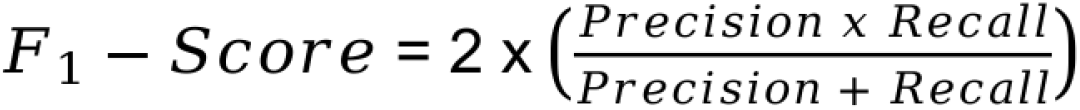

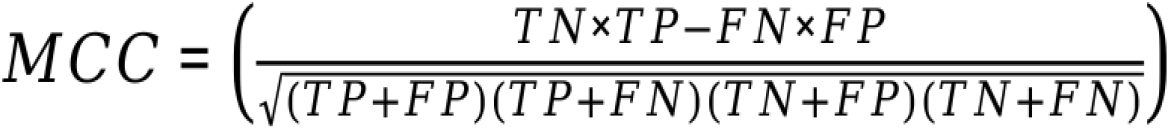

Where:

TP = True Positives, TN = True Negatives, FP = False Positives, FN = False Negatives, Acc = Accuracy.

A test was also conducted to determine if there was any significant over-fitting occurring in the Transformer-DenseNet final model. The gold standard method for detecting such over-fitting is through 10 times independent random training and testing trials, which compares the mean absolute error (MAE) between the training and testing performances. Each time, the dataset was randomly split in the ratio of 90:10, with the first part used for training and the second one for testing. Each time a new model was built from the scratch and evaluated on the corresponding test set. In addition, it was made sure that there was no overlap between the train and test sets to prevent any bias, and memory. This care has been taken for all the datasets taken in the present study.

**Full methods details are provided in the Supplementary Materials and Methods section. We strongly recommend readers to refer to this section for a comprehensive understanding of the methodological details employed in this study.**

## Results and discussion

### Learning on cumulative contextual information surrounding the anchoring prime motif helps identify TFBRs with high accuracy

The prime motif discovery (Detailed methods and results about which are given in the **Supplementary Information Results section**) helped in selecting the more appropriate contextual information and features. The motifs may occur significantly enriched in the binding data but by no means they are limited there only but also appear in the non-binding regions. The discovered motifs above worked as the point to zero upon to consider the potentially significant interaction spots across the DNA. Thus, it was imperative to assess their context and its contribution. Therefore, 75 bases flanking regions from both the ends of the motif were considered. Previously, it was found that such extent of the flanking regions around the potential interaction sites in nucleic acids capture the local environment contributory information effectively [37, 38, 47–50] as well as such regions were found important in determining the stationing of TFs through a localized search for right points to halt at [25]. The contextual information may come in the form of other co-occurring motifs, sequence and position-specific information, and structural/shape information which could work as strong discriminators against the negative and positive instances for TF binding. Considering the flanking regions around the motifs, three different datasets were constructed, Datasets “A”, “B”, and “C”, as described in the methods and related supplementary section. **Figure 2** illustrates how these datasets were built.

For building the models for 436 TFs and their binding preferences, 10 different combinations of various sequence representations were used. An assessment was made for each representation considered where the Dataset “A” was split into 70:30 ratio to form the train and test sets. This protocol worked as the ablation analysis to evaluate how each of these representations of the sequence was contributing towards the discrimination between the preferred binding and non-preferred binding regions through the transformer encoders (**Figure 4a**). The observed accuracy for dimeric representation was just 80.78% on Dataset “A”. This was followed by introduction of pentameric and heptameric sequence representations which returned the accuracy values of 85.05% and 86.23%, respectively, while covering a total of 156 and 154 words per sequence window, respectively. The ChIP/DAP-seq data does not retain strand information and complementary strands are also present almost equally and in most of the cases they too contribute in the binding. Considering the anchor motif’s counterpart from the complementary strand for the same binding region may boost the discriminating power further. Therefore, both strands were considered. By doing so, a significant improvement by ∼5% was noted for each of the individual representations. Yet, as can be seen here, individually all these representations displayed enough scope of improvement and needed information sharing with each other. Therefore, in the final stage, the datasets were formed having all the three representations of the sequences together. On these the transformers learned contextually along with the prime motifs with much higher amount of information sharing across the representations.

**Figure 4:**
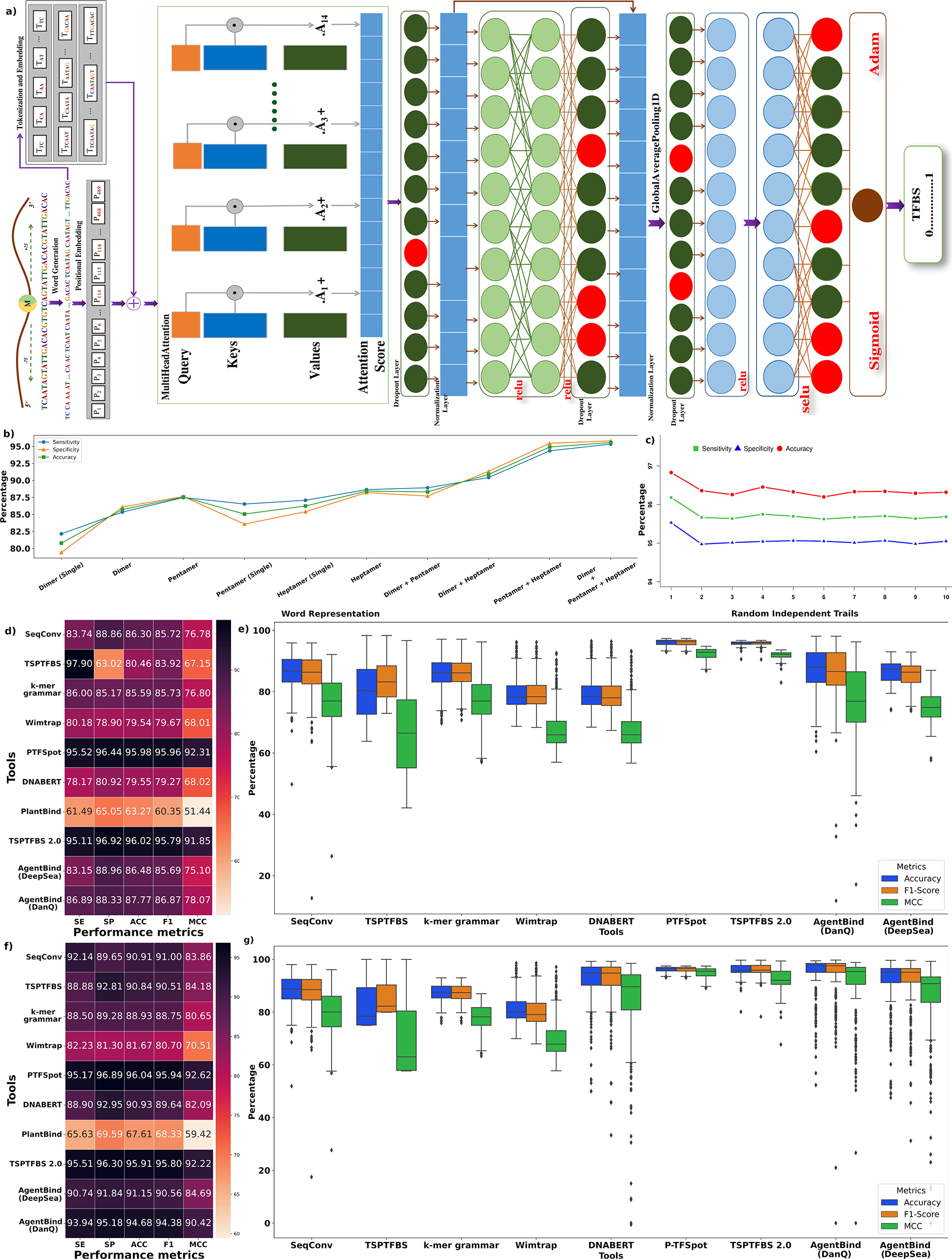
Transformer-only model’s implementation details. **(a)** This part utilizes DNA sequences based informations in terms of 7-mer, 5-mers and dimeric words while basing around (+-75 bp) the prime motif to detect the contextual information. **(b)** Ablation analysis for three main properties for discriminating between the negative and positive instances. These words representations appeared highly additive and complementary to each other as the performance increased substantially as they were combined together. (**c)** Ten-fold independent training-testing random trials on Dataset “B” depicts consistent performance of PTFSpot transformers. **(d)** Objective comparative benchmarking result on Dataset “B”. This datasets contained the TF originating from DAP-seq from the plant cistrome database. Here all the compared tools were train and tested on the Dataset “B”. **(e)** Performance dispersion plot on Dataset “B”. PTFSpot transformers consistently demonstrated minimal variance and distribution in its performance, maintaining a strong balance in identifying both positive and negative instances with a high level of precision. **(f)** Objective comparative benchmarking on Dataset ”C”. Here, all the compared tools were first trained and then tested Dataset ‘C’ and evaluated for their performance. This gave a clear view on the performance of each of the compared algorithms. **(g)** Performance dispersion plot on Dataset “C”. PTFSpot transformers consistently demonstrated high values across all the three performance metrics with least dispersion, affirming the algorithm’s robustness. From the plots it is clearly visible that for all these datasets and associated benchmarkings, PTFSpot consistently and significantly outperformed the compared tools for all the compared metrics (MCC values were converted to percentage representation for scaling purpose).

Combination of various representations of the sequences was done in a gradual manner in order to see their additive effect on the classification performance. These combinations of the words representations yielded a better result than using any single type words representations, as can be seen from **Figure 4b**. Complete details about word representations and performance can be found in **Supplementary Table 3 Sheet 1-3**. Details of the implementation of the optimized transformer are already given in the methods and associated supplementary section and **Figure 4a**.

10-fold random trials performance concurred with the above observed performance level and scored in the same range consistently. All of them achieved high quality ROC curves with high AUC values in the range of 0.9245 to 0.9561 (Dataset “A”) and 0.9562 to 0.9869 (Dataset “B”) while maintaining reasonable balance between specificity and sensitivity (**Figure 4c**; **Supplementary Figure 1; Supplementary Table 3 Sheet 4-5)**. To conduct an unbiased performance testing without any potential recollection of data instances, it was ensured that no overlap and redundancy existed across the data. The remarkable performance consistency ensured about the robustness of the raised transformer models and reliability of its all future results. It was evident that the Transformer effectively grasped both distant and nearby words associations, while acquiring knowledge through multiple hidden features.

### PTFSpot transformer models consistently surpassed all other compared TFBR finding tools

A highly comprehensive series of benchmarking studies was performed, where initially two different datasets, “B” and “C”, were used to evaluate the performance of the transformer models of PTFSpot with respect to nine different tools, representing most recent and different approaches of TFBR detection: AgentBind (DanQ, LSTM based), AgentBind (DeepSea, CNN based), *k*-mer grammar, Wimtrap, SeqConv, TSPTFBS, TSPTFBS 2.0, DNABERT, and PlantBind. The performance measure on the test set of Dataset “B” gave an idea how the compared algorithms in their existing form perform. The third dataset “C” was also used to carry out an objective comparative benchmarking, where each of the compared software was trained as well tested across a common dataset in order to fathom exactly how their learning algorithms differed in their comparative performance.

All these seven tools were trained and tested across Datasets “B” where PTFSpot-Transformers outperformed almost all of them, for all the performance metrics considered (**Figure 4d**). TSPTFBS 2.0 came very close to the performance of PTFSpot-Transformers with an average accuracy of 96.02% and MCC of 0.9185 (PTFSpot-Transformers average accuracy: 95.98%, average MCC: 0.9231). On the same Dataset “B”, the next best performing tool was AgentBind-DanQ (Avg accuracy: 87.77% and MCC: 0.7807) a very distant one in performance.

On Dataset “C”, PTFSpot transformers outperformed all the compared tools with significant margin with the similar level of performance (**Figure 4f**). PTFSpot transformers consistently demonstrated minimal variance in its performance, maintaining a strong balance in accurately identifying both positive and negative instances. This was evident through its high values across all the three performance metrics with least dispersion, affirming the algorithm’s robustness (**Figure 4e** & **g**). The full details and data for this benchmarking study are given in **Supplementary Table 3 Sheet 6-7**.

### The transcription factors vary across species and so do their binding regions preferences

One of the major calamities plant science has faced while exactly following the studies on humans is approaching TFs and DNA interactions in the similar fashion while overtly assuming binding sites conservation [31]. This is why in order to find TFs and their binding sites, *Arabidopsis* and TF specific models have been rampantly used, ending up with largely misleading results. All of them assume that the TFs and their binding preferences remain static and give no weight-age to their co-variability. While in actual, the transcription factors and their binding regions vary across the plant species [29, 30, 32].

As the TF structure changes, the binding sites also changes, and so does the preferred binding regions. Therefore, what works for one species, does not work for another until co-variability between TFs and their binding preferences is learned. And this is why most of the existing tools fail to work in cross-species manner, making them almost of no use. A study was done here to verify the same. Peak data of the common TFs were selected in *Arabidopsis thaliana* and *Zea mays* i.e., LHY1, MYB56, MYB62, MYB81, MYB88, and WRKY25. For both the species their corresponding prime motifs for the same TF were compared. It was observed that for the same TF different prime binding motifs existed. On comparing the same TF’s sequences and structures between the two species the following was observed:

1) The amino acid sequence identity for each TF was reasonably low between the two species, with lowest going up to 39% (LHY1). The amino acids sequence-based comparative details of each TF are given in **Supplementary Table 3 Sheet 8.**

2) The binding domain class of the compared TFs across the species were same but their amino acid compositions were different. For example, in the case of MYB88, both species contained two HTH DNA-binding domain, but an identity of only 33.3% was observed for the first domain and an identity of 43.2% was noted for the second domain when comparing both species (**Figure 5bI** & **5bII**).

3) The 3D structures of the same TF between the two species were compared and the RMSD difference between was found above 0.6 Å (**Figure 5**) [51]. This strongly suggested that the same TF varies significantly in its 3D structure when one goes across the various plant species. The superimposed structures for WRKY25 and MYB88 are given in **Figure 5aIII** and **aIV**, respectively. The details of 3D structure comparison of other TFs is given in **Supplementary Figure 2.**

**Figure 5:**
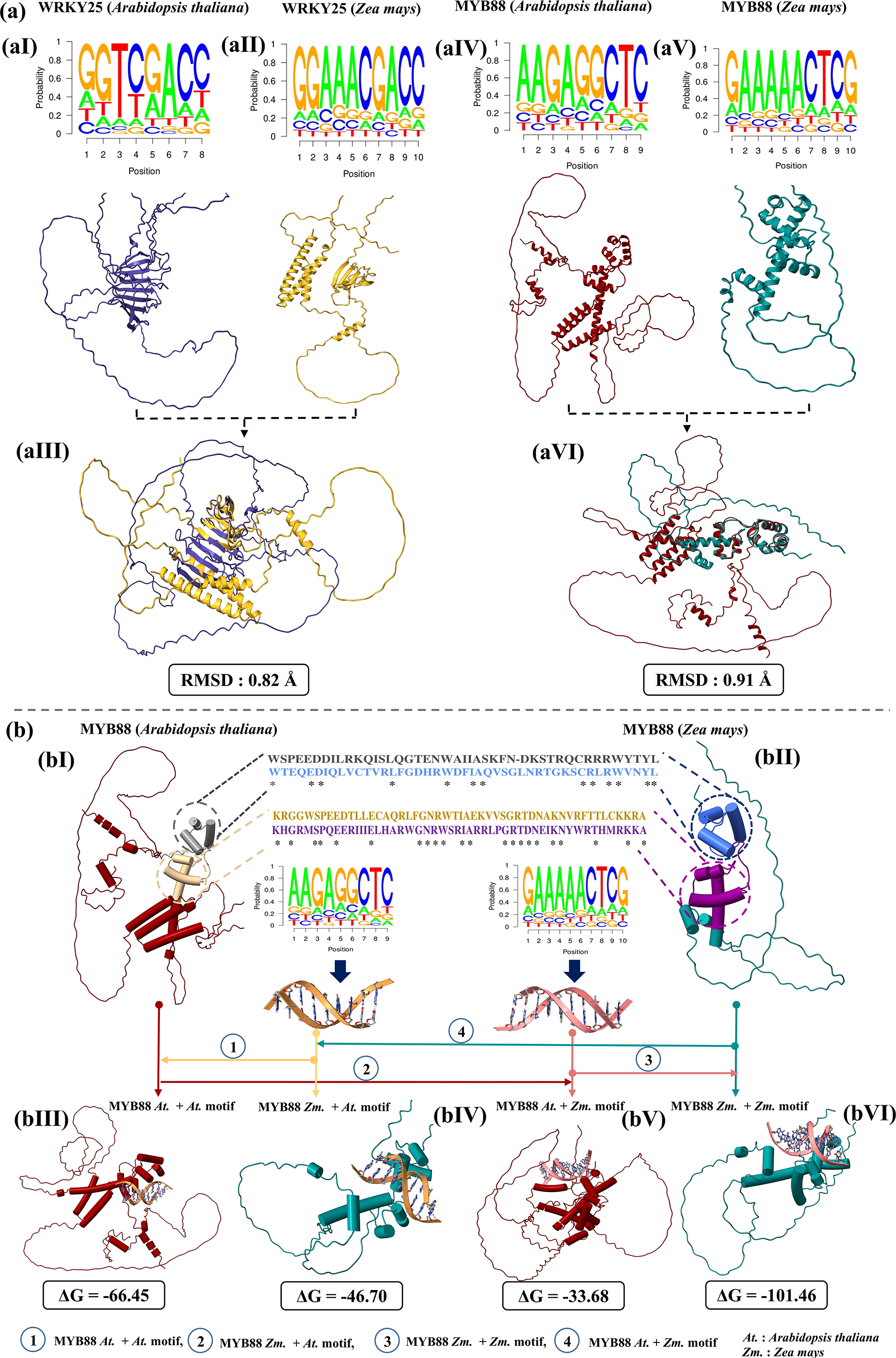
Co-variation in the structure of the transcription factor and the corresponding binding site across various species. **(a)** TF structure and its binding motif comparison across the species, **(aI)** prime binding motif of WRKY25 TF (*Arabidopsis thaliana*) and its 3D structure, **(aII)** prime binding motif of WRKY25 TF (*Zea mays*) and its 3D structure, **(aIII)** superimposed TF structures of *Arabidopsis thaliana* and *Zea mays*, with the structural differences measured in RMSD value. **(aIV)** prime binding motif of MYB88 TF (*Arabidopsis thaliana*) and its 3D structure, **(aV)** prime binding motif for WRKY25 TF (Zea mays) and its 3D structure, **(aVI)** superimposed TF structures for *Arabidopsis thaliana* and *Zea mays*, and corresponding structural difference in RMSD value. **(b)** Domain based comparison between *Arabidopsis thaliana* and *Zea mays* for MYB88, **(bI)** *Arabidopsis thaliana’s* MYB88’s has two domains. The first domain and its corresponding amino acids sequence are in grey color. The second domain and its corresponding amino acids sequence is shown in the tan color, **(bII)** *Zea mays* MYB88 too has two domains. The first domain and its corresponding amino acids sequence are in purple color. The second domain and its corresponding amino acids sequence is shown in maroon color, **(bIII and bVI)** The docking analysis shows the stability of the complexes when a TF was docked to its binding motif within the same species and to the one from another species for the same TF. It is clearly evident that the same binding motif can’t work across the species and it varies with species as well as the structure of the TF.

4) A TF’s binding affinity was higher to the prime motif within the same species than other species. As in **Figure 5**, taking the case of MYB 88, it was observed that when the TF of *Arabidopsis thaliana* was docked with its own prime motif, the binding affinity was −66.45 kcal/mol. However, when the *A*. *thaliana* MYB88 was docked with the prime motif for MYB88 of *Zea mays*, the binding affinity went much lower with −46.70 kcal/mol (**Figure 5bIII** & **bIV)**. The same procedure was applied for *Zea mays* MYB88 for its binding to its own prime motif and that of *A. thaliana,* and similar pattern was observed there also (**Figure 5bV** & **bVI**), clearly underlining that there is cross-species variability in TF binding preferences which is grossly neglected and leading to wrong study designs.

These important findings formed the foundation for the final form of PTFSpot as a universal TF-DNA interaction modeller which could work across any plant species and for even unseen TFs, while parallelly learning upon the structural variations and corresponding changes in binding region partner.

### Deep co-learning on TF sequence, structure, and corresponding DNA binding regions brings impeccably accurate universal model of TF-DNA interaction spots

During the real world application of cross-species identification of the TF binding regions, a huge performance gap exists, far below the acceptable limits. Some recent reports have highlighted the high degree of poor performance by a majority of the existing software tools for TFBR discovery during the process of their annotations where most of them end up reporting very high proportion of false positives [17, 18, 52]. Above, we have showed how variability in transcription factor structure and corresponding binding regions happen across the plants which none of the existing tools has attempted to learn. This becomes a major reason why the existing software pool does not work across the plant species.

As detailed in the methods sections about the architecture of PTFSpot implementation, a composite deep co-learning system was raised using Transformers and DenseNet which parallely learned upon the TF binding regions in DNA with sequence contexts and corresponding TF’s 3D structure and sequence. This model was trained and tested on Dataset “E” training and testing components, in 70:30 split ratio with absolutely no overlap and all redundancies removed. This co-learning system was trained using 41 TFs and their corresponding DAP-seq data. Each selected TF represented a single TF family and had highest binding data available among all the TFs for the respective family. The performance of the raised model achieved an excellent accuracy of 98.3% with a balanced sensitivity and specificity values of 97.56% and 99.04%, respectively (**Figure 6a**; **Supplementary Table 4 Sheet 1),** almost perfectly capturing the binding regions for every TF considered. The information sharing between the TF structure and binding region co-variability was so strong that when the model was assessed by removing the structural part, the accuracy dropped drastically to just 83.6%, significantly lower (p-value: 4.70e-24) than what was achieved above with co-learning on TF structure and corresponding binding data. The above raised model, unlike the existing ones, assumed all TFs falling into one hypothetical family whose structure and corresponding binding regions varied from one member to another, and this co-variability was learned to correctly identify the binding preferences for even those transcription factors on which it was not even trained.

**Figure 6:**
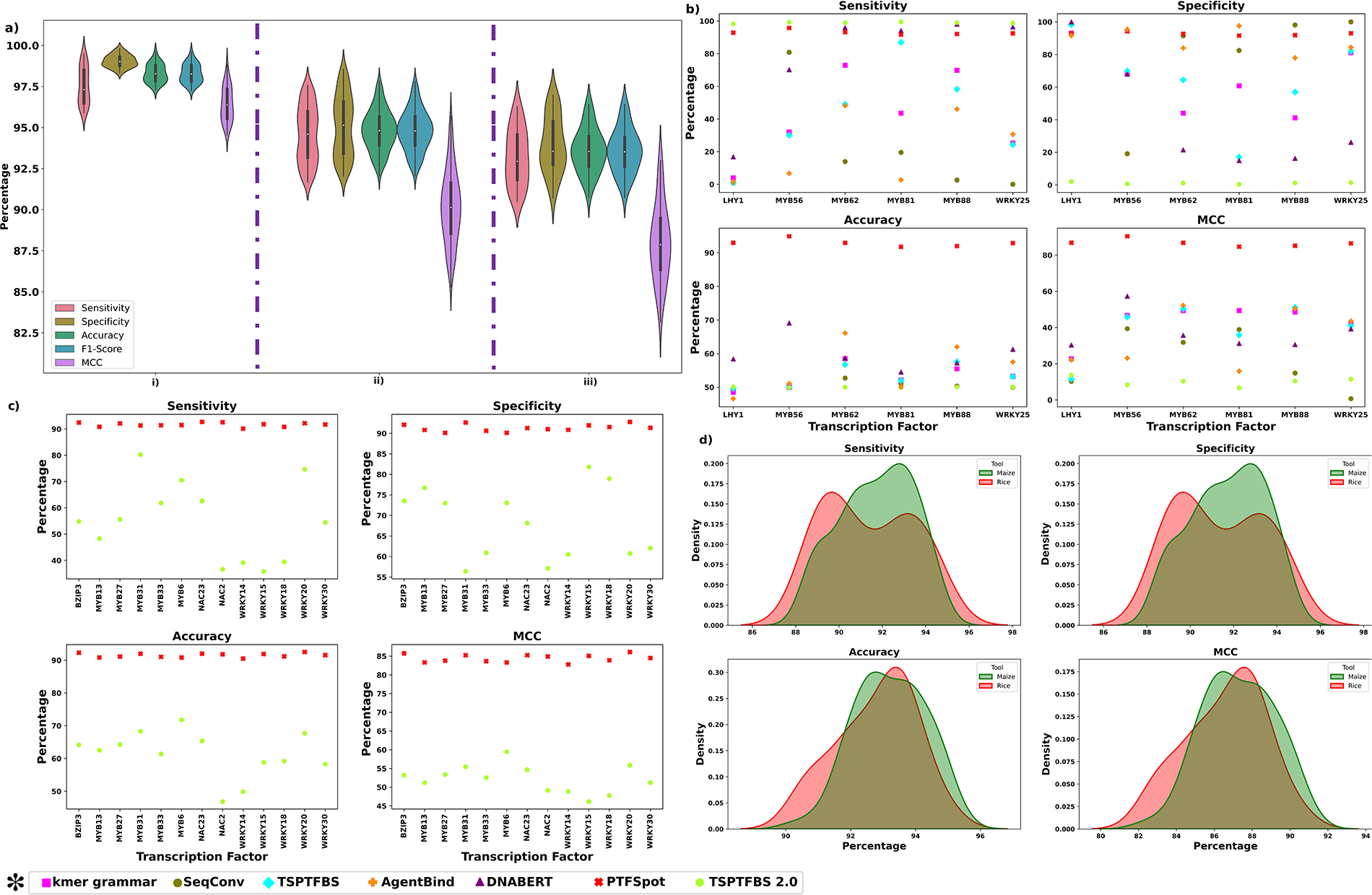
Performance and benchmarking of the universal model of PTFSpot. PTFSpot universal model was raised from 41 different TFs representing 41 different families, from *A. thaliana*. **a) (i)** The performance over the same 41 TF’s test set (Dataset “E”). **(ii)** The performance over the remain TF’s as test set from Dataset “E”. **(iii)** The complete dataset “D” (containing ChIP-seq TFs) worked as another test set. Performance on all of them was in the same range with exceptional accuracy. **(b)** Comparative benchmarking for the TFs whose models were available commonly among the compared tools. PTFSpot universal model outperformed them with huge leap for each TF. **(c)** Comparative benchmarking for the 13 wheat TFs (371,066 peak regions, equal number of negative instances following the same protocol as dataset “F”) whose models were available in TSPTFBS2.0 for comparison. PTFSpot universal model outperformed TSPTFBS2.0 for every compared TF by huge margins (p-value: 1.46e-05; Kruskal–Wallis test). Reason to select TSPTFBS2.0 for this comparison was that among the existing software tools, TSPTFBS2.0 was found the best performer. **(d)** PTFSpot universal model was raised using *A. thaliana* TFs. It was tested in trans-species manner across rice and maize. For both, it returned outstanding results, reinforcing itself as the solution for reliable cross-species discovery of TFBR.

The next important question was that how this co-learning system performed when introduced to different sets of transcription factors which had not even family representations in the above mentioned model which covered 41 TF families. The Dataset “D”, derived from Dataset “A”, had included 51 TF dataset. In the first part, we considered 21 TFs datasets exclusively. Each of these TFs represented a TF family which was never included in the training of the above mentioned co-learning model. Here, the co-learning model achieved an astonishing accuracy of 93.7% with balanced sensitivity and specificity values of 93.38% and 94.02%, respectively (**Supplementary Table 4 Sheet 2)**. A structural comparison based cluster analysis across all the 41 TFs in training and 21 TFs in the testing sets, both representing totally different TF family sets, was done. It was observed that performance observed for the most distant TF family in the test set was at par with the one with the closest distance with the members of the training set, and the overall performance across all the families was at the same level **(Supplementary Figure 3)**. Separate 10-folds training-testing runs were made to measure the performance consistency where all the runs scored at the similar level consistently. Also, the MAE for training was found to be 0.0348, while for testing it was 0.0362, resulting in a very small difference of only 0.0014. A t-test comparing the MAE values for the training and test sets yielded a highly insignificant result of ∼45%, much above the significance threshold of 5% or lower, further confirming the absence of any possibility of any significant over-fitting (**Supplementary Table 2 Sheet 2)**. This all concurred again that the model truly learned the co-variability between TF structure and its binding preferences with remarkable consistency and robustness.

Afterwards, we tested this model on the whole Dataset “D” where the total number of different TFs was 51. The average accuracy remained in the same range (93.6%) with very balanced sensitivity and specificity values of 93.2% and 94%, respectively (**Figure 6a**; **Supplementary Table 4 Sheet 2)**.

The application value of any such software is mainly when it is applicable universally, across various transcription factors (which PTFSpot qualified above) as well as across different species. More so when very frequently sequenced genomes of various plants are being released which essentially require TFBR annotations. Almost all of the existing software fail there.

The raised model’s last assessment was set to determine how well it performed to identify cross-species TF binding regions. For this task a new dataset, Dataset “F”, was employed completely as another test set. This dataset contained 117 TFs from *Zea mays* (93 TFs) and *Oryza sativa* (24 TFs), retrieved from GSE137972 (217 sample), GSE102920 (six sample), and ChIP-Hub with over 60 conditions. It was created solely for cross-species validation purpose for the model’s performance. In this study we included the most recent and advance tools like *k-mer* grammar, TSPTFBS, TSPTFBS 2.0, SeqConv, and Wimtrap to comparative benchmarking here. We also included a novel approach, AgentBind, based on context learning from the flanking regions which utilizes CNN to learn from the sequence patterns [15]. We looked for common TFs among different plant species to benchmark these tools for cross-species performance. In Dataset “F”, only six TFs from *Zea mays* (LHY1, MYB56, MYB62, MYB81, MYB88, WRKY25) were found for common ones which the existing software had any model developed (**Supplementary Table 4 Sheet 3)**. Thus, the comparative benchmarking was possible for only these six transcription factors for cross-species performance evaluation of the tools. For AgentBind, models for these TFs were raised using TF specific data used for PTFSpot while using AgentBind’s protocol.

Two levels benchmarking was performed. In the first level, seven tools, DNABERT, *k-mer* grammar, TSPTFBS, TSPTFBS 2.0, SeqConv, AgentBind, and Wimtrap, were considered and the six common TFs data were taken for evaluation along with PTFSpot universal model. PTFSpot universal model achieved an outstanding average accuracy of 92.9% ranging from 91.76% to 94.9%, clearly underlining its capability to accurately identify TFBR in cross-species manner. The performance observed for all the other tools was extremely poor, with none surpassing an average accuracy value of 60% (**Figure 6b**; **Supplementary Table 4 Sheet 3)**, reiterating the fact that none of the existing tools are suitable for any practical application like cross-species TFBR identification and genomic annotations. PTFSpot has emerged as a breakthrough solution to this situation.

An additional benchmarking exercise was done where TSPTFBS 2.0 performance was compared with PTFSpot for the DAP-seq binding data for 13 TFs (371,066 peak regions, equal number of negative instances following the same protocol as dataset “F”) from wheat [53]. These TFs were selected only because TSPTFBS 2.0 had models for only these TFs. TFPTFBS 2.0 was selected here because it was found as the best performing one among the existing pool of software. For this comparison also, PTFSpot significantly outperformed TSPTFBS 2.0 (p-value: 1.46e-05) with 30% performance lead over TSPTFBS 2.0 in terms of accuracy (31% lead in MCC) while attaining 91.48.% accuracy (84.43% MCC), clearly reiterating its impeccable performance in plant TF binding region discovery without any bounding to TF and species specific models, which none of the existing tools has attained so far.

In the next step, the above analysis was carried forward for the entire Dataset “F”, which covered data for 117 TFs from rice and maize. Here also, PTFSpot performed outstanding while attaining 93.58% average accuracy, MCC value of 0.87, and F1-score of 93.56%. **Figure 6d** provides further performance distribution illustration of PTFSpot for *O. sativa* and *Z. mays* transcription factors (**Supplementary Table 4 Sheet 4)**. All these series of validation and benchmarking studies proved that PTFSpot achieved a never seen before success in consistently and accurately identifying the binding sites for various families of transcription factor as well as across species due to its successful co-learning of the variability in structure and binding regions.

To give a short glimpse of the kind of impact PTFSpot could have due to its capabilities to detect co-variability between TF structure and its binding preferences, we extended our above described case MYB88 example. MYB88 is reported to influence PIN7 gene involved in auxin efflux and transport associated with plant development [54] by binding at five locations in the promoter of PIN7. When TSPTFBS2.0 was run to detect the same in *A. thaliana,* it could not report any binding site for MYB88. However, PTFSpot detected all of them there. The homologous gene for PIN7 in Maize is zmPIN1c [55] whose promoter was also scanned for MYB88 binding. This gene displays ∼60% identity between Maize and *Arabidopsis* and exhibit a remarkable variability despite retaining its function. As already showed in the section above, even the structure of MYB88 has drifted a lot from *Arabidopsis* to Maize. When *Arabidopsis* specific Transformer-only PTFSpot model of MYB88 was run to scan for its binding regions in the promoter of zmPIN1c in Maize, nothing was found. TSPTFBS2.0 also reported nothing for MYB88 there. However, when the universal model based PTFSpot was run, it detected two binding locations for MYB88 there. To validate any possible regulatory role of MYB88 in transcriptional regulation of zmPIN1c in Maize, we performed gene expression correlation analysis, utilizing data from nine experimental condition available at Maize Expression Atlas database (European Bioinformatics Institute: [https://www.ebi.ac.uk/gxa/home]. Remarkably, a very strong Pearson correlation coefficient of 0.97 was observed, indicating a very high possibility of regulation of zmPIN1c by MYB88 in maize. Further details of this analysis are presented in the **Supplementary Table 5-6**. This small demonstration highlights the kind of deep impact PTFSpot may have in unraveling the regulatory systems of plants while breaking several age long bottlenecks.

## Conclusion

The present work brings a revolutionary new approach, PTFSpot, which learns from the co-variability between binding protein structure and its binding regions without requiring to be specific for any particular transcription factor or its family specific model. It can accurately identify the binding regions for any given transcription factor belonging to any family across any plant genome and can work for even any novel and never reported before transcription factors and genomes with same level of accuracy. With this, the present work is expected to drastically change the scenario of plant regulatory research as well as may cause extensive cutting of cost incurred on experiments to detect TF binding regions across a genome.

## Acknowledgments

The work was carried out under the aegis of The Himalayan Centre for High-throughput Computational Biology (HiCHiCoB), a BIC supported by DBT, Govt. of India. SG and VK are thankful to DBT, India for financial support as project associateship. UB is thankful for DBT JRF fellowship. Jyoti is thankful for CSIR-UGC SRF fellowship. The authors are thankful to Ritu for her inputs on DenseNet. This MS has CSIR-IHBT MSID 5451.

## Author’s contributions

SG carried out the major parts of this study and developed the web-server and standalone versions of PTFSpot. VK carried out the structural analyses. Jyoti, UB, and VK carried out the benchmarking study. RS conceptualized, designed, analyzed, and supervised the entire study. SG, VK, UB, Jyoti, and RS wrote the MS.

## Software and Data availability

All the secondary data used in the present study were publicly available and their due references and sources have been provided in **Supplementary Tables 1–6**. The software has also been made available at GitHub at https://github.com/SCBB-LAB/PTFSpot as well as a webserver at https://scbb.ihbt.res.in/PTFSpot/ (all related datasets in the study are hosted here).

