## Supplementary Information for "PTFSpot: Deep co-learning on transcription factors and their binding regions attains impeccable universality in plants"

### Materials and Methods

#### Data retrieval

For *Arabidopsis thaliana*, ChIP-seq and DAP-seq peak data were collected for 441 TFs from PlantPAN3.0 [1] and Plant Cistrome Database [2]. While Plant Cistrome Database has 387 TFs cover 44 families with over 568 files of two different origins, i.e., i) DAP-seq and ii) ampDAP-seq, while data from PlantPAN3.0 has 54 TFs cover 25 families with over 293 conditions. **Figure 1c** presents the number of collected TFs across 56 different families. All the data were collected in the form of coordinates. The peak data varied from 16 (HBI) to 52,997 (EPR1) peaks. Five TFs data were initially removed because there was not enough peak data availability. Therefore, only TFs with more than 400 distinct binding peaks were considered. HBI (16), TOC1 (191), GATA19 (202), DDF2 (326), and AP2 (329) were the five TFs with the fewest initial peaks. The remaining 436 TFs data contained a total of 5,753,198 distinct peaks. With all this, 5,753,198 peaks, ranging from FHY3 (444) to EPR1 (52,997), were accessible for the study. From the genome, initial coordinate information was extracted into sequences. The source and type of collected peak data are given in **Supplementary Table 2 Sheet 1**.

#### Identification of motif seed candidates

In the initial phase of the motif search, the peak regions of ChIP-seq and DAP-seq were scanned to identify the most prevalent 6-mer seeds with at least 70% identity. Reasoning behind this was the fact that several previously described TF motifs considered at least six bases [1-4] while functional conservation is usually expected within the range of 70% identity. To satisfy a lower bound for such motifs, a *k*-mer with six bases and at least 70% identity would look for all 6-mers spectra that match each other. To determine the enrichment status of the *k*-mers (seeds), all possible *k*-mers were generated for each such sequence which were repeatedly and simultaneously searched through all the TF-bound regions indicated in the targets. Previously, we had introduced a similar motif and seed discovery algorithm to detect RBP-binding sites [5]. We implemented the same motif and seed discovery algorithm to detect the motif seed candidates for the TFs.

**Anchoring with the significant seeds**

The peak data was assessed for the shared *k*-mer seeds and their relatives with at least 70% identity. Further analysis was done for the statistical importance of the seed candidates. For any TF, the best representative seeds were those which had a representation rate of at least 75% across the data. The occurrence probabilities of each *k*-mer and its relatives were computed using the random dataset to determine their null model distribution. A random dataset was generated from the genome by following peak data length distribution. Binomial test was done for significant enrichment analysis between binding and random data. Null model probabilities were computed using the randomly selected regions from DNA regions not reflected in the ChIP/DAP-seq data. All those *k*-mers which qualified the binomial test at 1% significance level were considered for the further analysis as the significant ones. These significantly enriched *k*-mers were the initial seeds for constructing the final motifs. Bidirectionally extension of the seeds was done iteratively by adding one nucleotide on each side at each step, followed by peak data search with at least 70% identity match. The final motifs were chosen based on satisfying of both criteria: the motif candidate had to exhibit at least 75% abundance in the peak data at 1% significance level as well as all matching instances displayed at least 70% identity. Such final motif which was most dominant tagged as the "*prime motif*". In addition, a TF could recognize both the strands of the DNA at a given position. Thus, the reverse complementary of the prime motif was also looked to strengthen the information. Positive and negative datasets for a specific TF were built simultaneously using the input matrices (position weight matrices (PWM)) for the forward and reverse-complement motifs.

**Comparison with experimentally reported motifs**

We retrieved 197 TFs from JASPAR [3] that overlapped with our set of 441 TFs. A total of 201 motifs were reported for these 197 TFs. The length of the retrieved motifs ranged from 8-24 bases. To maintain similar motif search approach for fare comparison, these motifs were scanned similarly to those found using the PTFSpot approach. Experimentally verified motifs from Plant Cistrome Database were also considered in addition to the JASPAR motifs. This way a total of 589 motifs were retrieved. Similar searches were also done for these motifs. The relevance of the resemblance between the experimentally reported motif and the corresponding matching motif from PTFSpot was assessed using TOMTOM, a part of MEME suite [6].

**Datasets creation**

The corresponding peak data sequences were transformed into instances of positive data sets once the motifs were anchored for each TF in the provided peaks data. The two ends of the found motifs were extended by -75 and +75 bases in both the directions to produce positive dataset instances for each TF while considering the genomic sequences after mapping the peak data. All sequences in any given sample and corresponding TF had the same length. Due to the motif's repeated occurrences inside a single sequence, the overall number of instances exceeded the peak data for most of the TFs.

Many genomic sites resemble interaction motifs found in the bound cases, but they do not interact in actual. In such scenario, taking random sequences into account does not significantly boost the aim of discrimination. Therefore, one needs to be extremely careful while creating a sound negative dataset. Considering those genomic regions which display the regions similar to the prime motifs found enriched in ChIP-seq TF binding data but never appeared in ChIP-seq data, become very strong negative dataset candidates. Such negative instances capture the natural scenario of binding preferences where over importance of binding motif is downplayed.

Therefore, we removed all those regions from the genome which were reflected in the peak data, ensuring that the remaining regions never reflected in the peak data and were fit for negative instances selections. Next, they were searched for the prime and reverse complementary motifs of the TF similar to the positive data cases. Similarly, 75 bases flanks were considered while capturing the contextual information with more discrimination power [7]. Positive and negative instances were pooled for each TF in a 1:1 ratio without any overlap.

The dataset where the positive instances were derived from ChIP-seq studies has been called Dataset “A” throughout this study. On the other hand, the dataset whose positive instances were derived from DAP-seq has been named as Dataset “B”. Additionally, a different dataset containing 387 TFs binding data was formed using data from the Plant Cistrome Database. They too were from DAP-seq studies. This dataset was also utilized by TSPTFBS [8] and TSPTFBS 2.0 [9] for their study. The creation of positive data required the conversion of TFs peak data into FASTA format. We used a large pool of 1,89,799 background sequences provided by TSPTFBS, randomly chosen from the intergenic regions of the TAIR10 *Arabidopsis thaliana* genome, to collect the negative data. The ratio of positive to negative data was kept as 1:1. This dataset has been named as Dataset “C” in the current study.

Further, two more datasets “D” and “E” were created from datasets “A” and “B”, respectively for the development of universal and generalized model for TF-DNA interactions and its testing. The only difference between datasets “A” and “D” and datasets “B” and “E”, was that 3D structures of the associated TFs were also considered in the development of these Datasets “D” and “E”. 3D structure of the associated TF were derived with the help of Alphafold2 [10]. For every positive and negative instance, 3D structure of that associated TF was taken as the input. For Dataset “D”, the total number of TFs was 51. For Dataset "E", the total number of TFs was 325. Both the datasets contained TFs for *Arabidopsis thaliana*. Dataset "D" was kept aside purely for independent testing purpose only and Dataset "E" was taken into account for model training and testing.

Besides Dataset "D" and Dataset "E", one more independent test dataset, known as Dataset "F" was also generated. This dataset was constructed to extend the examination of the generalization of the raised model for cross-species TFBS identification. This dataset contained 117 TFs from two species *Zea mays* (93 TF) and *Oryza sativa* (24 TF). This dataset has instances from GSE137972 (217 sample), GSE102920 (six sample), and ChIP-Hub [11] with over 60 conditions. A total of 2,278,620 ChIP-seq peaks were collected. For the positive binding instances of a given TF, we extended the peak region by mapping to its respective genome. For the negative instances of each TF, we randomly selected the sequences with the same sequence length and numbers as the positive instances after excluding the positive instances information for the considered species. It was ensured that any overlapping regions between negative and positive instances for each of the TFs were removed and finally these non-overlap regions were used as negative dataset instances for each TF. Dataset "F" is also another independent dataset kept aside in entirety and purely for testing purpose alone. For all the datasets every possible overlap and redundancies (full or partial) were screened out.

**Word representations of sequence data**

Each sequence can be considered as a collection of independent words. For instance, the dimeric words AT, TT, TG, GG, GC, CA, and AG represent the sequence ATTGGCAG. An alphabet of the four bases allows creating 16 unique 2-mer words, 1024 unique 5-mer words, and 16,384 unique 7-mer words. Every occurrence of the dinucleotide, pentamer, and heptamer sequences were transformed for the verbal representation of the sequences in an overlapping window.

Dinucleotides, pentamers, and heptamers provide information on composition and base stacking, shape, and regional motif information, respectively [5, 12, 13]. The input vector had a maximum length of 469 words and a sequence length of 159-162 bases. Word representations as dinucleotide, pentameric, and heptameric sequences were independent. Therefore, distinct integers were used as a token for each unique word. The tokenized version of the sequence were converted further into numeric vectors and matrices, known as embedding. The transformer encoder fed the tokenized and embedded sequences as input to train and assess the models. The TensorFlow (Keras) tokenizer class was used to implement the tokenization procedure.

**Implementation of the Transformers**

A deep-learning (DL) strategy was used to construct TFBS discovery models in plants using tokenized and embedded vectors. The transformer-encoder's multi-headed attention mechanism generates reliable contextual correlations. They are placed into hidden space vectors, which the model's output layer categorizes.

The sequences from the previous step's tokenized representation were fed into the input layer of the Transformer system. To create an output matrix formatted as samples, sequence length, and embedding size, embedding converted each word's token into a word vector whose length equals the embedded size. There were "***n***" words in each sequence of length "***l***" (sentence size). The process of embedding words into vectors is represented as following:

Consider a text-based input sentence, **S** = { $x_1$ , $x_2$ , $x_3$ ……, $x_n$ }, where $x_n$ represents the $n^{th}$ word in the sentence. Each word in a sentence is transformed into a *d-dimensional* vector, where *d* (28) represents the dimension of the sequence element representations in the transformer model, commonly referred to as the model dimension, whose elements ( $I_d$ ) carry optimized numeric weights, resulting in high level embedding vector, $x_n$ =[ $I_{n1}$,$I_{n2}$……,$I_{nd}$ ].

Accordingly, the text-based sentence can be represented as an embedded words matrix, $M \in \mathbb{R}^{n\ x\ d}$

$$M = \begin{bmatrix} x_1 \\ x_2 \\ x_3 \\ .\ . \\ x_n \end{bmatrix} \begin{bmatrix} I_{11}, I_{12}, \ldots, I_{1d} \\ I_{21}, I_{22}, \ldots, I_{2d} \\ I_{31}, I_{32}, \ldots, I_{3d} \\ .\ .\ .\ .\ .\ .\ .\ .\ .\ .\ .\ .\ . \\ I_{n1}, I_{n2}, \ldots, I_{nd} \end{bmatrix}$$

where each row corresponds to a word in "***S***". This matrix has dimensions of *n* x *d*, indicating the number of words in the sentence by the dimension used to represent each word in the embedding vector.

Each word in the matrix "***M***" is also combined with its corresponding positional encoding, "***P***". The positional encoding "***P***" shares the same dimension "***d***" as the word embedding vector, $x_n$, resulting in M' = M + P, where ***P*** = [ $p_1, p_2, \ldots, p_d$ ]. The calculation of sinusoid positional encoding "***P***" are determined using the following equations:

$$P_{(i,\ 2m)} = sin\left(\frac{i}{10000^{2m/d_{model}}}\right)$$

$$P_{(i,\ 2m+1)} = cos\left(\frac{i}{10000^{2m/d_{model}}}\right)$$

where, "***i***" denotes the position of token and "***m***" stands for the encoding dimension.

Position encoding generates a matrix with a similar shape to be added to the embedding matrix. The shape of the matrix (samples, sequence length, embedding size) remains consistent throughout the Transformer and is eventually reshaped by the final output layers. The input embedding layer feeds its outputs into the subsequent layer, while the output embedding layer does the same for the next layer (encoder layer).

The encoder processes the input through a multi-head attention layer. The multi-head attention applied to

$M' \in \mathbb{R}^d$

is computed in the following manner. The MultiHead result is obtained by concatenating h individual attention heads and

$(head_1, head_2, ..., head_h)$

multiplying them by a weight matrix $W^O$ .

$$MultiHead = Concat\ (head_1, head_2, ..., head_h)W^Q$$

Each attention head (***head$_i$***) calculates the attention mechanism with input matrices Q, K, and V. These input matrices are derived from "***M'***" using optimizable weight matrices $W^Q$ , $W^K$ , and $W^V$ .

$$head_i = Attention\ (Q, K, V)$$

where, Q = M'. $W^Q$ ,

K = M'. $W^K$ ,

V = M'. $W^V$

The attention operation itself is defined by:

$$Attention(Q, K, V) = Softmax\left(\frac{QK^T}{\sqrt{d_k}}\right) \bullet V$$

This entire process is applied to input matrices Q, K, and V. Finally, the ***h*** attention head outputs are concatenated and further transformed using an output weight matrix $W^O$ .

The same steps were repeated and their individual attention vectors were concatenated and passed to the transformer-encoder's input feed-forward network block for additional processing. To lessen overfitting, the output of the multi-head attention layer was supplied to a dropout layer, followed by a normalization layer. After normalization, the output goes into two fully connected feed-forward layers, which converges to the subsequent dropout layer, followed by the normalization layer. A GlobalAveragePooling1D layer was then implemented, which feeds its results to the third dropout layer of the model after the previous layer's input was collected via two fully connected feed-forward layer. Various activation functions from the available activation functions were examined for the layers. The final hidden layer passes its output to fourth dropout layer. A one-node classification layer using the sigmoid activation function with binary cross-entropy loss was implemented. The weights and learning rates were adjusted in this step with the help of the "Adam" optimizer. **Figure 4a** depicts the operation of the implemented transformers system.

**Optimization of the Transformer system**

The transformer system was optimized using Bayesian optimization. The transformer encoders had a multi-head attention layer, where 14 self-attention heads were found to work best. The input sequence was padded if the length was shorter or longer than 160 bases to ensure a constant size of the input matrix. The output from the multi-head attention layer was passed into the dropout layer with a dropout fraction of 0.15.

Later, this result was normalized by another layer called the normalization layer. This layer was followed by a third layer called feed-forward layer with 14 nodes, followed by another dropout layer with a dropout fraction of 0.15. A feed-forward layer with 14 nodes, followed by another normalization layer, followed the above one. The result of the normalization layer was passed to the GlobalAveragePooling1D layer, followed by a third dropout layer with a dropout fraction of 0.25. Next to this, the pooled feature maps were passed to fully connected layers. The performance of the transformer was evaluated for several hidden layers where two hidden layers were found performing the best. The number of nodes in two hidden layers was set for the model, and different activation functions for the layers were explored among the available activation functions. Finally, two dense layers were configured, each with 40 and 16 hidden nodes with SELU and RELU activation functions, respectively. The last hidden layer was connected to the fourth dropout layer of the stack with a dropout fraction of 0.25, which finally passed its results to a single-node result classification layer based on a sigmoid activation function.

A binary cross-entropy loss function was used to calculate the loss and "Adam" optimizer was used to adjust the weights and learning rates. The learning rate of the optimizer was set to 0.001, and the model was trained using 25 epochs and batch size of 16. Since the problem in this study was not translation but classification, decoders were not needed. The number of encoders and performance was investigated, and it was found that increasing the encoder layers did not show a significantly large change in efficiency and only decreased after the sixth encoder layer. Thus, we went with one encoder layer to keep it lighter with the same performance.

The output of the transformers returns probability scores for each input sequence indicating the confidence of each case as the presence or absence of TFBS in the given sequence. The final hyperparameters of the output layer of the optimized model were: {"Activation function": sigmoid, "Loss function": binary cross-entropy, "Optimizer": Adam}. Related information from the optimization to the final model is listed in **Supplementary Table 2 Sheet 3** and illustrated in

**Supplementary Figure 4**. The resulting final model was saved in Hierarchical Data Format 5 (HDF5). Since the whole system is implemented in TensorFlow, the HDF5 format provided the model graph definition and weights to the TensorFlow structure and saved the model for classification. **Figure 3** shows the detailed workflow and the implemented transformer architecture part in it.

**Performance evaluation for the transformer part**

According to standard practice, the datasets "A" and "B" were divided into train (70%) and test datasets (30%). The developed model was tested on the 30% intact and completely untouched test portion. Four categories of performance confusion matrix namely true positives (TP), false negatives (FN), false positives (FP) and true negatives (TN) were evaluated. The performance of the built transformer models was evaluated using performance metrics such as sensitivity, specificity, accuracy, F1 score, and Matthews correlation coefficient (MCC) [14].

Performance measures were done using the following equations:

$$Sensitivity\ (Sn) = \left(\frac{TP}{TP + FN}\right)$$

$$Specificity\ (Sp) = \left(\frac{TN}{TN + FP}\right)$$

$$Acc = \left(\frac{TN+TP}{TN+TP+ FN+FP}\right)$$

$$F_1 - Score = 2 \text{ x } \left(\frac{Precision \text{ x } Recall}{Precision + Recall}\right)$$

$$MCC = \left(\frac{TN \times TP - FN \times FP}{\sqrt{(TP+FP)(TP+FN)(TN+FP)(TN+FN)}}\right)$$

$$AUC = \int_0^1 Pr\,[TP](v)\,dv$$

Where:

TP = True Positives, TN = True Negatives, FP = False Positives, FN = False Negatives, Acc = Accuracy, AUC = Area Under Curve.

In addition, 10 times independent random training and testing trials were used to assess the consistency of the model and its performance. Each time, the dataset was randomly split in the ratio of 90:10, with the first part used for training and the second one for testing. Each time a new model was built from the scratch. For each of these models, the above mentioned performance metrics were calculated. In addition, it was made sure that there was no overlap between the train and test sets to prevent any bias, and memory. This care has been taken for all the datasets taken in the present study.

**Comparative benchmarking**

The developed approach, PTFSpot, was comparatively benchmarked with nine different tools that reflect most advanced current approaches for TFBS detection. The compared tools were AgentBind (DanQ, LSTM based), AgentBind (DeepSea, CNN based), *k*-mer grammar (bag-of-*k*-mers and vector-of-*k*mers), Wimtrap (XGBoost), SeqConv (Convolution Neural Nets (CNN)), TSPTFBS (CNN), TSPTFBS 2.0 (DenseNet), PlantBind (composite CNN, Long Short Term Memory (LSTM), and Attention mechanism) and DNABERT (Bidirectional Encoder Representations from Transformers (BERT)). Additionally, the benchmarking took into account two distinct datasets (Dataset “B” and “C”) to conduct a completely unbiased evaluation of the performance of these tools. The reasoning to build Dataset “C” was to use the same set of data which was used by the compared tools to report their performance [8, 9, 15-18]. Details on the datasets are already covered in the methods section.

**Structure and binding motif analysis across species**

We further studied the counterion binding of TFs to the target DNA sequences across the species. We downloaded the available common TFs (*Arabidopsis thaliana* and *Zea mays*) peak data from PCBase and Plant Cistrome Database. The 3D structure of the TFs was modeled using AlphaFold2 [10]. AlphaFold2 generated five models for each TF, but we selected the top-ranked model for our subsequent analyses.

ScanProsite (ExPASy) (http://prosite.expas.org) was used to confirm each modeled protein's functional domain and the number and name of amino acid residues found in the pocket of the active site. PTFSpot was applied to identify motifs for all these TFs across the species. Afterwards, we did a comparative investigation into 3D protein structure with their corresponding motif for all the common TFs taken. Comparative study of each TF was done on the following bases: 1) based on the protein sequence, 2) based on the functional domain, 3) based on their respective 3D structure, and 4) the binding affinity of TF across the species.

**Validation of prime motif and transcription factor interactions using molecular docking and simulation**

For this study, we randomly selected four TFs (AT566940, CEJ1, WRKY75, and VRN1) and their identified prime binding sites. The prime binding sites identified through PTFSpot algorithm for above mentioned four TFs were taken as flexible molecules. Each of them were studied for binding with the identified prime motif region and other regions around. Four more regions were taken for each of the TF. These regions were selected from sequences to evaluate the impact of flanking regions around and the prime motif region. In doing so, for each such region was removed and binding analysis was performed. Region ‘A’, represented the prime motif region, whereas region ‘B’ and ‘D’, were overlapping regions with the prime motif in both the directions. Region ‘C’ and ‘E’ were taken from the start and end of the sequence for every considered TFBS, respectively (**Supplementary Figure 5c & 5d**). The pyDockDNA server [19] was used to calculate the binding affinity and selectivity for these regions (A, B, C, D, and E) with its TF protein. From each docked complex (with regions A, B, C, D, and E) with the lowest docking energy, the optimal complex structure was chosen for each. The strength of the binding region and target interaction was determined by the structural qualities of the target. The GROMACS package [20] was used to run the molecular dynamics simulation (MD) for a protein-DNA complex structure utilizing the amber99sb force field. Each complex was solvated in a cubic box with 1.2 nm distance between the complex and the solvated box edges. To neutralize the system's charge, sodium ($Na^+$) and chloride ($Cl^-$) ions were added, and the energy was minimized using the steepest descent algorithm. A 2 fs time step was used in the simulation of each complex structure. All simulations of the complexes were undertaken for a 50 ns to verify the robustness of the results. Chimera [21], PyMOL [22] and Xmgrace software were used to analyze the results.

**Transformer-DenseNet system for cross-species identification of the binding regions**

Initially, only Transformer-based model was implemented to identify TF binding regions for every individual TF. Any such model fails to perform well when run across other species for the same TF family. This happens due to variation in the TF structure and its binding regions within species. As discussed above, protein 3D structure plays an important role in determining binding motifs. We incorporated TF’s structure information for every corresponding binding region from ChIP-seq/DAP-seq data. 3D protein structure based DenseNet [23] and the above developed Transformer model which takes sequences as input, were intertwined through a bi-modal architecture implementation to raise a universal model to identify TF-DNA interactions in global and cross-species manner.

The DenseNet part of PTFSpot takes atom-wise X, Y, Z coordinates of the 3D structure of a protein as the input. We normalized the coordinates by dividing it by the highest absolute value noted for the corresponding axis. This normalization ensured transformed and uniform values in range of -1 to 1. This coordinates based matrix was the input with 300×24×3 dimensions to a convolution layer. If the length of some protein was found shorter, it was padded for the empty columns with value of zero. Size of 300 covers the amino acid positions in a window, for each of which corresponding 24 atoms exist with every atom mapped to three-dimensional space in terms of normalized X, Y, Z coordinates. Those amino acids which have less than 24 atoms in their conformation were padded for the empty columns with value of zero. The 24 elements vector size to represents the amino acids was considered after an analysis of more than 400 TFs where we found that the TF amino acids had maximum of 24 atoms in it. Thus, each vector position represented a particular atom number.

**Building the DenseNet architecture**

Input Layer: The input to the DenseNet is the 3D tensor representing the protein structure. The dimensions was 300×24×3 (300 amino acids, 24 atoms per amino acid, three X, Y, Z space coordinates for each atom).

DenseNet: We opted for DenseNet in our study due to its inherent advantages in mitigating the vanishing-gradient issue, enhancing the propagation of features, promoting feature reuse, and significantly minimizing the parameter count. Each layer receives a “collective knowledge” from all the preceding layers. Since each layer receives feature maps from all the preceding layers, this leads to higher computational and memory efficiency. The architecture of the DenseNet model employed in our study is composed of one convolution layer with 32 convolution filters (kernel size = 3), one batch normalization and 2D maxpooling pooling layer (stride = 2), ten dense blocks and nine transition layers (**Figure 3**). A total of 121 layers were implemented in DenseNet.

Within each dense block, ‘ $m_0$ ’ represents the input to the convolutional network, consisting of ‘ $l$ ’ layers. Consequently, the ‘ $l^{th}$ ’ layer receives its input from the feature maps of all previous layers, which is represented as:

$$m_l = \mathrm{H}_l\left(\left[m_0, \mathrm{m}_1, \ldots\ldots, \mathrm{m}_{l-1}\right]\right)$$

Here, $\left[m_0, \mathrm{m}_1, \ldots\ldots, \mathrm{m}_{l-1}\right]$ denotes the concatenation of feature maps generated in layers 0 to $l-1$. $H_l$ (.) represents a non-linear transformation, which can be described as a composite operation consisting of batch normalization (BN), followed by a rectified linear unit (ReLU) function, and a 3x3 convolution (kernel size = 3). To facilitate down-sampling, the network architecture incorporates transition layers between each dense block. These transition layers include batch normalization, followed by a 3x3 convolution layer, maxpooling2D, batch normalization, 3x3 convolution layer, and conclude with a dropout layer. The growth rate, denoted by the hyperparameter ‘***k’*** in DenseNet, plays a pivotal role in the architecture's remarkable performance. DenseNet can achieve impressive results even with a smaller growth rate, primarily due to its unique design that treats feature maps as a global state of the network. In each layer, ‘***k’*** feature maps are contributed to this global state, and the total number of input feature maps (denoted as $F_l$) at the ‘***l-th’*** layer is calculated as:

$$F_l = k_0 + k \bullet (l-1)$$

The term ‘ $k_0$ ’ represents the number of channels in the input layer. After the final transition layer, the initial input, which was of shape 300x24x3, had been down-sampled to a final size of 1x1x64. Subsequently, the output of DenseNet was flattened and concatenated with that of transformers for the final classification steps. In the second part of the bi-modal architecture transformers were implemented with sequence as input constructed as described in above section on transformers. Afterwards, output passes on the GlobalAveragePooling1D layer which it passes to dropout and then to one hidden layer. From both parts (DenseNet and Transformers), outputs from the previous layer were concatenated and passed onto the batch normalization, dropout, dense, and dropout layers sequentially. A sigmoid activation function based single node classification layer was used with binary cross entropy loss function to calculate the loss. “Adam” optimizer was used at this point to adjust the weights and learning rates. The batch size was set to 64 and the number of epochs was set to six. Further details of this module is provided in the **Figure 3**.

To train and test this bi-modal Transformer-DenseNet system, two different datasets were created from (Datasets “D” and “E” derived from the datasets “A” and “B”, respectively). Details are already covered in the datasets section above. Considering the TF protein structure and the corresponding binding region together while co-learning on them captures the relationship between protein structures and binding sites and variations in them. As mentioned in the methods section, Dataset “D” was kept for independent testing purpose only and Dataset “E” was taken into account for model training and testing. Data were split in the ratio of 70:30 on which the bi-modal system was trained and tested. Besides Dataset “D” and “E”, one more independent test dataset, known as Dataset “F” was also generated. This dataset contained 117 TFs from two species *Zea mays* (93 TF) and *Oryza sativa* (24 TF). Dataset “F” is considered as the independent test dataset to evaluate the performance of the developed Transformer-DenseNet based bi-modal deep learner for cross-species

TF binding regions identification in plants. Details on all datasets is already covered in the datasets related section in the methods section.

### Results

#### Most of the TF binding regions display a prime binding motif

It was observed that each of the TFs considered in this study had at least one such prime motif which covered at least 75% of the binding data. The prime motifs ranged from 9 to 12 bases. Full details on the prime motifs and their abundance for each studied TF is given in **Supplementary Table 2 Sheet 4-5.**

The TFs formed 48 distinct clusters (**Supplementary Figure 6 and 7**) out of similarity among their prime motifs, an observation consistent with the earlier works which had observed that TFs from the same family had may display similar binding motifs **[3, 24]**. Even the members from vast and functionally diverse bZIP and NAC families displayed close proximity and shared similar prime motifs, concurring previous observations **[25, 26]**.

The motifs reported in the present study were compared with the experimentally reported motifs using TOMTOM. Many of the motifs found in this study matched with the experimentally reported motifs (**Supplementary Table 2 Sheet 6-8**). It was also observed that several of these experimentally reported motifs were not the prime motif reported here but matched with other lower ranked motifs which either co-occurred with the prime motifs or were exclusively present, covering a comparatively lesser amount of ChIP/DAP-seq binding data than the prime motifs reported in the present study. It may be due to that fact the prime motifs reported here are through a huge binding data of ChIP/DAP-seq with much bigger experimental space than previously reported motifs, influencing their relative ranking. Their occurrence in the data varied from 3.42% to 99% while the prime motifs reported in this study mostly covered at least 75% of the bound data (**Supplementary Table 2 Sheet 6-8**). **Supplementary Figure 8** provides a snapshot of comparison between motifs reported in this study and their experimentally reported counterparts. A standard structural and molecular dynamics based validation study on the identified prime motifs reconfirmed them important for binding to the TF and supported the TF:DNA complex.

**Structural and molecular dynamics analyses support the prime motifs as important one for TF binding**

The docking results revealed that the identified prime motif regions were showing the highest binding affinity with their corresponding TFs than the other regions. The detailed information of docking score of all TFs are given in **Supplementary Table 2 Sheet 9.**

Comparative analysis of Root Mean Squared Deviation (RMSD) trajectories of five different TFs-DNA complexes from each sequence partner to the interacting TFs were studied for stability. The trajectory was measured at 300K for 50 ns. The complexes with the prime motifs were found to be much more stable than the other regions. The method section provides the details on various regions considered. For example, in case of DOF (ATDOF5), comparative analysis of RMSD value for the Apo (unbound form of protein) DOF, the value ranged from 0.5 to 2.7 nm while for the prime motif consisting complex A, the value ranged from 0.5 to 2.4 nm and stabilized at 2 nm. Similarly, RMSD value for the complex B, the values ranged from 0.5 to 2.3 nm and stabilized at 2.2 nm, which was lesser stable. Complex C and D had RMSD values ranging from 0.1 to 4 nm and 0.5 to 2.2 nm and got stabilized at 3.9 nm and 2.1 nm, respectively. For the complex E, RMSD showed deviation from 0.1 to 3.8 nm and settled at 3.5 nm (**Supplementary Figure 5j**). In all the five complexes, the sequence with the prime motif (complex A) was found to be more stable when compared to the others in the dynamic environment. Moreover, the second sharpest impact was observed for the overlapping and immediately flanking regions around the prime motif. As one moves away from the prime motif region, the impact on binding diminishes. The identified prime motifs provided structural stability to the considered TF-DNA complexes (**Supplementary Figure 5e-i**). A similar pattern was observed for other three TFs taken in this study (**Supplementary Figure 9**). The docked complex and its binding affinity values are provided in **Supplementary Figure 5**.

**Supplementary Table 1. List of some published tools for Transcription factor binding site identification**

| S. No. | Software | Algorithm | Encoding scheme | Biological relevance | Dataset | Species | Year | Webserver (W)/ Standalone (S) |
|---|---|---|---|---|---|---|---|---|
| **1** | **KmerHMM [27]** | HMM | kmer | Model the dependence between adjacent nucleotide positions | PBM | Human | 2013 | W/S |
| **2** | **GkmSVM [28]** | SVM | gapped-kmer | Detection and modulation of functional sequence elements in regulatory DNA | ChIP-seq | Human | 2014 | S |
| **3** | **pPromotif [29]** | Probabilistic modeling | position weight matrix, conservation index (Ci Value), and inter-nucleotide dependence | Plant transcrip-tion factor binding sites | AGRIS database and TBFS annotations in GenBank entries | *Arabidopsis thaliana* | 2014 | S |
| **4** | **DeepSEA [30]** | CNN | One-hot encoding | Identify the noncoding-variant effects *de novo* from sequence on chromatin | ChIP-seq | Human | 2015 | S |
| **5** | **DeepBind [31]** | CNN | One-hot encode | Nucleic acid binding site | ChIP-seq | Human | 2015 | S |

| | | | | | | | | |
|---|---|---|---|---|---|---|---|---|
| | | | | prediction and can discover new patterns even when the locations of patterns within sequences are unkonwn | | | | |
| **6** | **Basset [32]** | CNN | One-hot encode | Learn the complex code of DNA accessibility across many cell types and annotate every mutation in the genome with its influence on present accessibility and latent potential for accessibility | DNase-seq | Human | 2016 | S |
| **7** | **DanQ [33]** | CNN and Bi-LSTM | One-hot encode | Identify non-coding function *de novo* from sequence which can have enormous benefit for both basic science and translational research | ChIP-seq, DNase-seq | Human | 2016 | S |
| **8** | **LS-GKM [34]** | SVM | Gapped kmer | Identify and detect the regulatory vocabulary encoded in functional DNA elements, and significantly contribute to understand gene regulation. | ChIP-seq | Human | 2016 | S |
| **9** | **DeeperBind [35]** | CNN+LSTM | One-hot encoding | Model the positional dynamics of probe sequences and hence reckons with the contributions made by | PBMs | Human | 2017 | S |

| | | | | | | | | |
|---|---|---|---|---|---|---|---|---|
| | | | | individual sub-regions in DNA sequences | | | | |
| 10 | **DeepSNR [36]** | CNN and DeepCNN | One-hot encoding | Identify TF binding location at Single Nucleotide Resolution *de novo* from DNA sequence and learns the dependencies between nucleotides at different positions within the binding site description | ChIP-exonuclease | Human | 2018 | S |
| 11 | **TFimpute [37]** | CNN | One-hot encoding | Identify cell-specific TF binding | ChIP-seq | Human | 2017 | S |
| 12 | **KEGRU [38]** | Bi-GRU | *k*-mer embedding | Capture complex context information from the k-mer sequence | ChIP-seq | Human | 2018 | S |
| 13 | **DeFine [39]** | CNN | One-hot encoding | Identifies cell type-specific functional impact of abundant non-coding variants including SNPs and indels. Also identifies the causal functional non-coding variants from disease-associated variants in GWAS | ChIP-seq | Human | 2018 | W/S |
| 14 | **K-mer grammar [17]** | Logistic regression | *k*-mers | Framework to exploit characteristic chromatin contexts and sequence organization to classify regulatory regions based on sequence features - *k*-mers | ChIP-seq, Mnase-seq | *Zea mays* | 2019 | S |

| | | | | | | | | |
|---|---|---|---|---|---|---|---|---|
| **15** | **FactorNet [40]** | CNN+BiLSTM | One-hot encoding | Identify cell type-specific transcription factor binding by leveraging signal data, such as DNase I cleavage | ChIP-seq | Human | 2019 | S |
| **16** | **DESSO [41]** | CNN | One-hot encoding | Identify motifs and identify TFBSs in both sequence and regional DNA shape features | ChIP-seq | Human | 2019 | S |
| **17** | **DeepRAM [42]** | CNN/ RNN | One-hot/*k*-mer embedding | Uses Different architectures using CNNs or RNNs to identify DNA/RNA sequence binding specificity | ChIP-seq | Human | 2019 | S |
| **18** | **WSCNNLSTM [43]** | Multi-instance learning and hybrid neural network | *k*-mer embedding | Identify *in-vivo* protein-DNA binding | ChIP-seq | Human | 2019 | S |
| **19** | **AgentBind [44]** | CNN + BiLSTM | One-hot encoding | Score the importance of context sequences | ChIP-seq | Human | 2021 | S |
| **20** | **DLBSS [45]** | CNN/ RNN | One-hot encoding and shape features | Identify TF-DNA binding preference using input DNA sequences and shape properties | PBM | Human | 2021 | S |
| **21** | **TbiNet [46]** | CNN, Bi-LSTM, and Attention mechanism | One-hot encoding | Identify transcription factor binding sites | ChIP-seq | Human | 2020 | S |
| **22** | **FCNA [47]** | CNN | One-hot encoding | Accurately identify TF-DNA binding motifs | ChIP-seq | Human | 2021 | S |

| | | | | | | | | |
|---|---|---|---|---|---|---|---|---|
| | | | | across different cell lines and infer indirect TF-DNA bindings | | | | |
| 23 | **BPNet [48]** | CNN | One-hot encoding | Discove relevant motifs and syntax rules underlying the cis-regulatory code | ChIP–nexus | Human | 2021 | S |
| 24 | **SAResNet [49]** | Self-attention mechanism+residual network | One-hot encoding | Identify DNA-protein binding and learning of the long-range dependencies from the DNA sequence | ChIP-seq | Human | 2021 | S |
| 25 | **SeqConv [15]** | CNN | One-hot encoding | Identify more precise TF-DNA interaction regions in plants | ChIP-seq | *Zea mays* | 2021 | S |
| 26 | **TSPTFBS [8]** | CNN | One-hot encoding | TFBS prediction in plants | DAP-seq | *Arabidopsis thaliana* | 2021 | S |
| 27 | **DNABERT [18]** | BERT | *k*-mer encoding | Enables direct visualization of nucleotide-level importance and semantic relationship within input sequences for better interpretability and accurate identification of conserved sequence motifs and functional genetic variant candidates. | ChIP-seq | Human | 2021 | S |
| 28 | **D-AEDNet [50]** | Deep Attentive Encoder-Decoder Neural Network | One-hot encoding | Identify the location of TFs–DNA binding sites in DNA sequences at base-pair level by leveraging the nucleotide position information | ChIP-exo and ChIP-seq | Human | 2021 | S |

| | | | | | | | | |
|---|---|---|---|---|---|---|---|---|
| **29** | **DeepGRN [51]** | CNN+Bi-LSTM+Attention mechanism | One-hot encoding | Automatically and effectively predict transcription factor binding sites | ChIP-seq, DNase-Seq | Human | 2021 | S |
| **30** | **Wimtrap [16]** | XGBoost | PWM | Identify condition- or organ-specific *cis*-regulatory elements and TF gene targets, with a great flexibility regarding the input data | ChIP-seq | *Arabidopsis thaliana* | 2022 | S |
| **31** | **PlantBind [52]** | CNN+Bi-LSTM | One-hot encoding | Identify potential TFBSs of multiple TFs simultaneously | ChIP-seq | *Arabidopsis thaliana* | 2022 | S |
| **32** | **MAResNet [53]** | Top-down and bottom-up attentation mechanism+residual network | One-hot encoding | Identify transcription factor binding sites in DNA sequences | ChIP-seq | Human | 2022 | S |
| **33** | **FCNsignal [54]** | Encoder + Decoder + Skip Architecture | One-hot encoding | (i)Identify base-resolution signals of binding regions, (ii) discriminating binding or non-binding regions, (iii) locating TF-DNA binding regions, and (iv) Identify binding motifs. | ChIP-seq and ATAC-seq | Human | 2022 | - |
| **34** | **TSPTFBS 2.0 [9]** | DenseNet CNN | One-hot encoding | TFBS prediction in plants | DAP-seq | *Arabidopsis thaliana* | 2023 | S |

**HMM: Hidden Markov Model; SVM: Support Vector Machine; CNN: Convolutional neural network; LSTM: Long Short Term Memory; GRU: Gated Recurrent Unit; RNN: Recurrent Neural NetwOrk; BERT: Bidirectional Encoder Representations from Transformers

**Supplementary Table 5: Binding region of MYB88 on PIN7 and zmPIN1c in *Arabidopsis thaliana* and *Zea mays* respectively.** This table presents the results of the PTFSpot analysis detailing MYB88 binding regions identified within the promoter sequence of PIN7 in *Arabidopsis* and zmPIN1c in *Zea mays*. **(a)** depicts the binding regions of MYB88 on PIN7 in *Arabidopsis*, revealing a total of five distinct binding regions characterized by their respective start and end positions within the gene promoter. **(b)** illustrates the binding regions of MYB88 on zmPIN1c in *Zea mays*, identifying two binding sites along with their start and end positions within the promoter sequence.

| **a) Binding region of MYB88 on PIN7 in *Arabidopsis thaliana*** | | | |
|---|---|---|---|
| **S.No.** | **Start** | **End** | **Binding region in promoter sequence** |
| **1** | 113 | 273 | AGTTTAAAAAACAAAATATATGAAGCAGAATTTCAAAATCATTAA<br>GATAAAATTATTCGCGGGATCAGTGATAGAATCTCAATGTAATTTT<br>TTCAACTTTCGTTTGCATCGACACAAAATCAGTGATAGAATCTTA<br>ATGTAATTTTTTCAAAAAAGTTTA |
| **2** | 521 | 681 | CCCATGTTATTAAGACAATCCCCTTCTATCTGTTTATGTGACGACT<br>ATACCGATAAAGATTTCTGATTACGTAAGACATTTTCGATAATACC<br>AAGAGATAAAATATCTTCTCTCGTCGTATATATTGAAATACATGGT<br>GCACGGAGTTCATTATAGCTTT |
| **3** | 908 | 1068 | GATCCTCCACCCACCAATATTTAATCTTAGGGATGTACAATTTATC<br>TGTAATATTATAGTACCTACAATTACTTGTTTCTACTTCAACCAAA<br>AATATGAATTTTCATAGTAGAATTAAATTCAAAAACCAAAAAAAA<br>ACTTAATGTTGGTCCATAACTGG |
| **4** | 1201 | 1361 | TCGTTCATCGCAAAATCAGGCGTAATAGTCGGAGACTCTATTTAA<br>GATATTAACAGCAACACAACAAATTATGATCTTTTTACGTCACCTA<br>AAAAAGGGTCATTTTCAATCGTTTTCTATATTGGAGATTTGAGAA<br>CATACATACCTAGAAATAAAAGAG |
| **5** | 1487 | 1647 | TTTTTATTTTTCTATATATAATTTGTGTTTACATCACAAAGCTTACT<br>AAAACGACGACACCGCGTTTTGAGTGCAAAGCGCCTAACCCACT<br>TCTGAATGGTAACCAATGCTCTGAATGGTAACTAGAGCTTTATCA<br>AGTCGACGTACTTGTCTAAACAGC |
| **b) Binding region of MYB88 on zmPIN1c in *Zea mays*** | | | |
| **1** | 560 | 720 | GGTTGGTATAAGTAAAATATAGTTTTAAGCATCATGGTTTACTCAA<br>AAATATACCTGATCATAGGCCACCCGACTCGACCAATGCAACAGC<br>CAGGTGTGGGCCATAATTTTTGACCCGCAATAATTCACGAGCGGG<br>TACTAGCCATGAATTGTTGACCCA |
| **2** | 817 | 977 | GCCATGCTAAATGCCCCACGACCCGACATTGACTCGACCCAACG<br>TTTAACAGGTCTACTCAAAATCGTGGTATCCTTTTGAGTTTTGAA<br>ACTTCACTCAGGACCTTTGTTTTCTTTTATTCTCTCTACAAATACT<br>TCTTTTTCAGTTTTGAAACTTCACT |

**Supplementary Table 6: Gene expression correlation analysis: Expression profiles of MYB88 and zmPIN1c in various tissues.**

| | Expression value in TPM | | | | | | | | |
|---|---|---|---|---|---|---|---|---|---|
| **Gene** | **anther** | **ear** | **embryo** | **leaf-base** | **leaf-middle** | **leaf-tip** | **root** | **shoot** | **tassel** |
| **zmPIN1c** | 4 | 3 | 2 | 3 | 4 | 0.5 | 21 | 3 | 4 |
| **MYB88** | 12 | 0.5 | 0 | 0 | 2 | 2 | 87 | 11 | 0.5 |
| **Number of samples: 18** | **Pearson corr** | **0.97** | | | | | | | |

### Supplementary data

**Supplementary Table 1**. List of some published tools for Transcription factor binding site identification

**Supplementary Table 2**: This supplementary file holds various data. Sheet 1 has details for various ChIP-Seq experiments and DAP-seq experiments for the studied TFs. The second sheet provides details on ten fold random independent trails details. Sheet 3 provides details on Hyperparameters for the transformers of PTFSpot. Sheets 4 and 5 cover the prime motifs details for Datasets "A" and "B", respectively. Sheets 6-8 cover details on experimentally reported motifs. Sheet 9 covers the details of docking study.

**Supplementary Table 3**: The supplementary file contains details for nine different studies done in the MS which cover i) Impact of words representation on the transformers performance. ii) sheets 2-3 cover performance of PTFSpot on the datasets "A" and "B". iii) sheets 4-5 cover details on 10 fold independent random trials for datasets "A" and "B". iv) sheets 6-7 cover performance benchmarking studies on datasets "B" and "C". v) Sheet 8-9: Details of the study showing the variations in sequence and domains of the transcription factors when compared between *A. thaliana* and *Z. mays*.

**Supplementary Table 4**: Performance evaluation details for the universal model of PTFSpot across various experimentally validated datasets.

**Supplementary Table 5:** Binding region of MYB88 on PIN7 and zmPIN1c in *Arabidopsis thaliana* and *Zea mays* respectively. This table presents the results of the PTFSpot analysis detailing MYB88 binding regions identified within the promoter sequence of PIN7 in *Arabidopsis* and zmPIN1c in *Zea mays*. **(a)** depicts the binding regions of MYB88 on PIN7 in *Arabidopsis*, revealing a total of five distinct binding regions characterized by their respective start and end positions within the gene promoter. **(b)** illustrates the binding regions of MYB88 on zmPIN1c in *Zea mays*, identifying two binding sites along with their start and end positions within the promoter sequence.

**Supplementary Table 6:** Gene expression correlation analysis: Expression profiles of MYB88 and zmPIN1c in various tissues.
