## Supplementary Figures for "PTFSpot: Deep co-learning on transcription factors and their binding regions attains impeccable universality in plants"

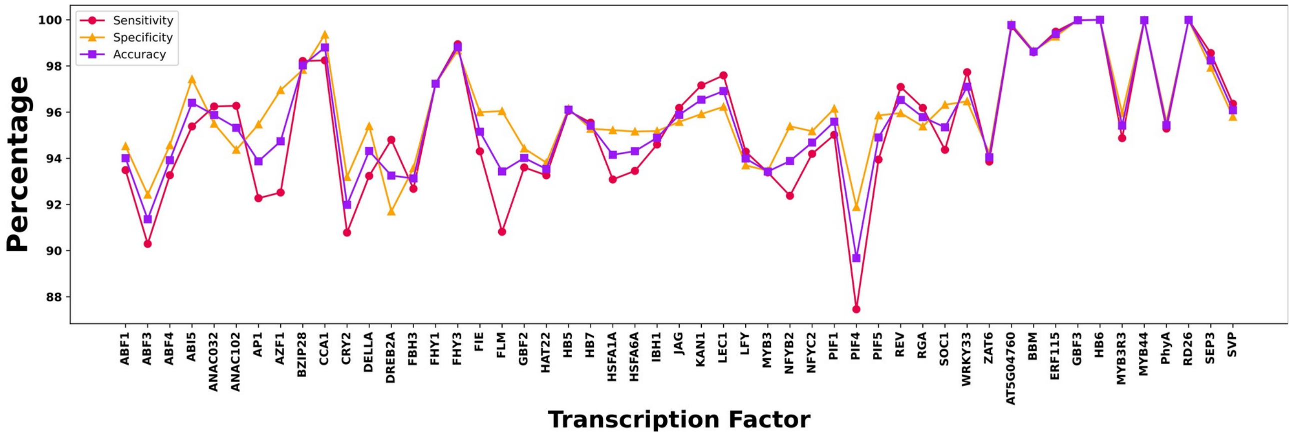


**Supplementary Figure 1: Performance results for PTFSpot transformer for Datasets "A".** The developed approach demonstrated a consistently strong and dependable performance across the Dataset "A".

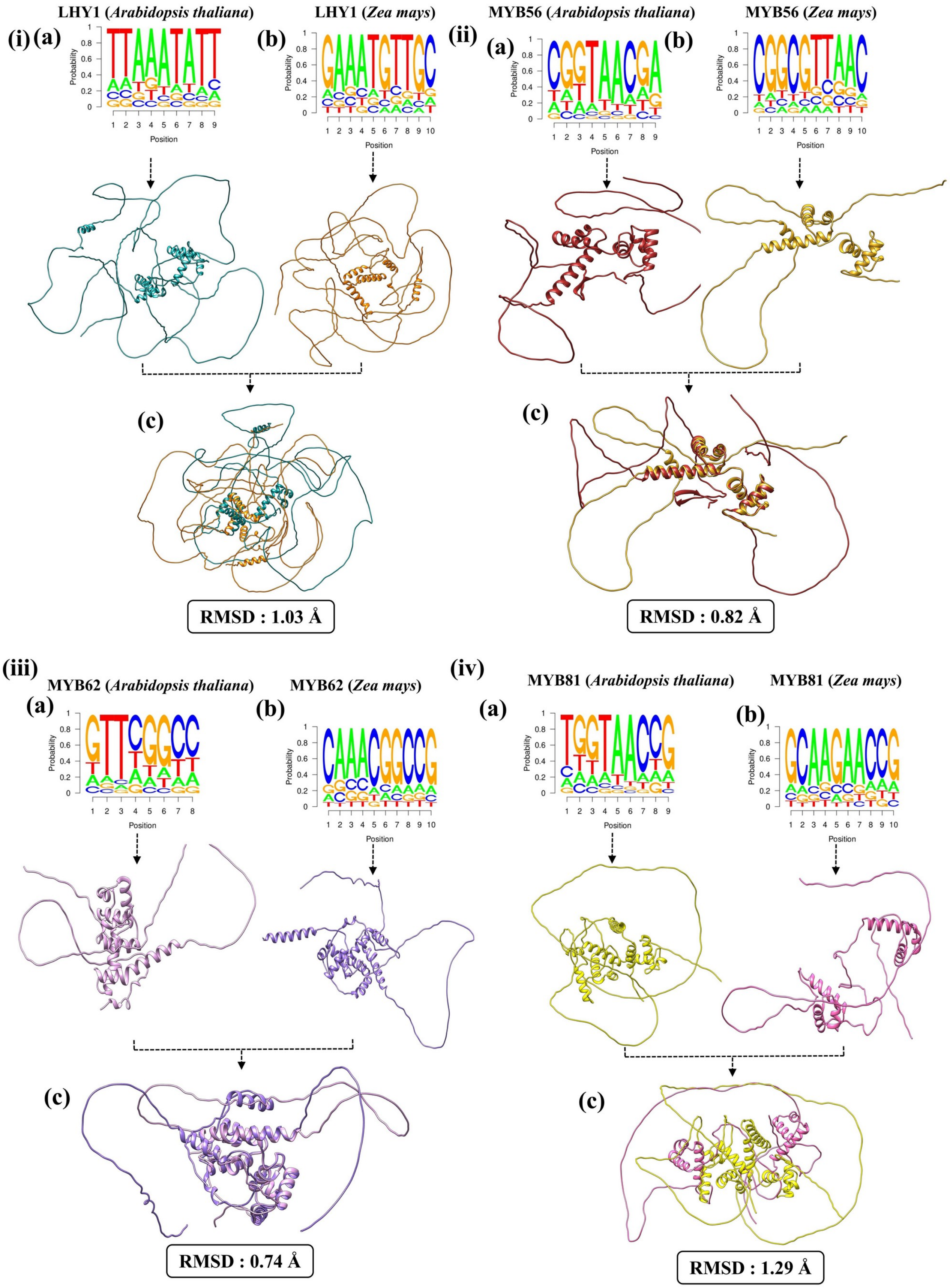

(i)
LHY1 (Arabidopsis thaliana)
LHY1 (Zea mays)
(a)
(b)
(c)
RMSD : 1.03 Å
(ii)
MYB56 (Arabidopsis thaliana)
MYB56 (Zea mays)
(a)
(b)
(c)
RMSD : 0.82 Å
(iii)
MYB62 (Arabidopsis thaliana)
MYB62 (Zea mays)
(a)
(b)
(c)
RMSD : 0.74 Å
(iv)
MYB81 (Arabidopsis thaliana)
MYB81 (Zea mays)
(a)
(b)
(c)
RMSD : 1.29 Å
Probability
Position

**Supplementary Figure 2: 3D structure based co-variation in transcription factors and their binding sites across the species (*A. thaliana* vs *Z. mays*). (i)** The prime binding motifs of LHY1 and their corresponding superimposed TF structures with the RMSD value indicating the structural variations. (ii) The prime binding motifs for MYB56 and their superimposed TF structures, with the structural differences expressed as the RMSD value. (iii) The prime binding motifs for MYB62, along with the comparison of their superimposed TF structures and structural differences expressed as the RMSD value. (iv) The prime binding motif for MYB81 and their superimposed TF structures with the structural differences measured in terms of the RMSD value. Further details of this figure is covered in Figure 5.

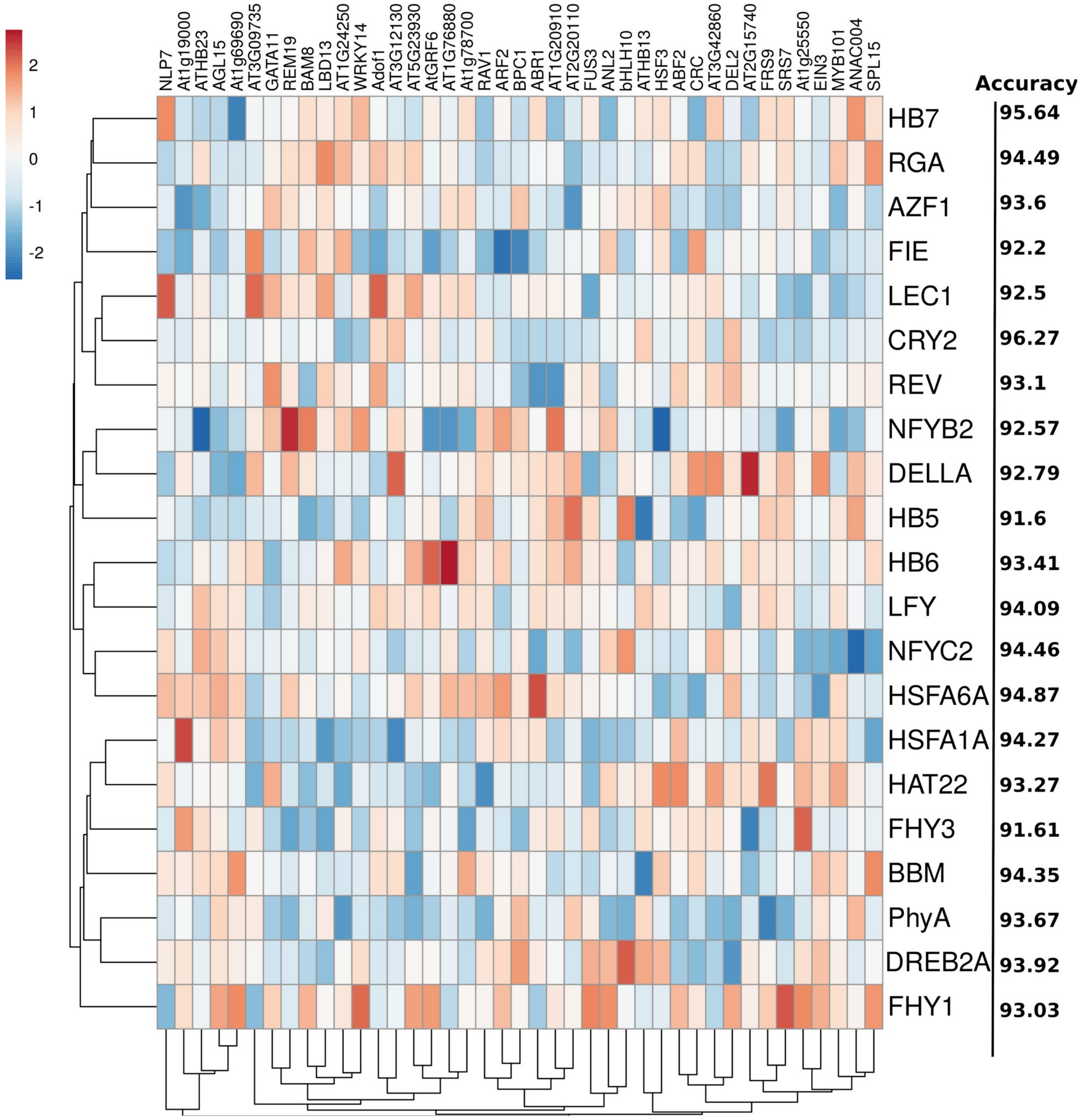


**Supplementary Figure 3: Structural similarity comparison between representative transcription factors for various TF families from training and testing datasets and their associated accuracy values.** This figure illustrates the structural similarity clustering of 41 transcription factors (TFs) used for training and 21 TFs used for testing, along with the accuracy achieved by the PTFSpot algorithm for the testing set TF's binding regions discovery. The heatmap depicts the structural divergence between the TFs, with red indicating higher dissimilarity and blue indicating higher similarity. The TF FHY1 exhibited the highest structural divergence from all the training data TF members, with an average RMSD of 1.25Å, suggesting a substantial structural difference. Despite this, FHY1 was detected with 93.03% accuracy by PTFSpot, which falls within the claimed performance range and is considered a very good value. In contrast, the PhyA TF showed the highest structural similarity to the training data, with an average RMSD of 0.9Å. Notably, the identification accuracy for PhyA (93.67%) was in a similar range as FHY1. In overall, for all the studied TFs, despite varying structural similarities, the performance range remained the same, clearly indicating that PTFSpot has learned the covariability between structure of the TF and its binding regions  rather than simply memorizing the training cases.

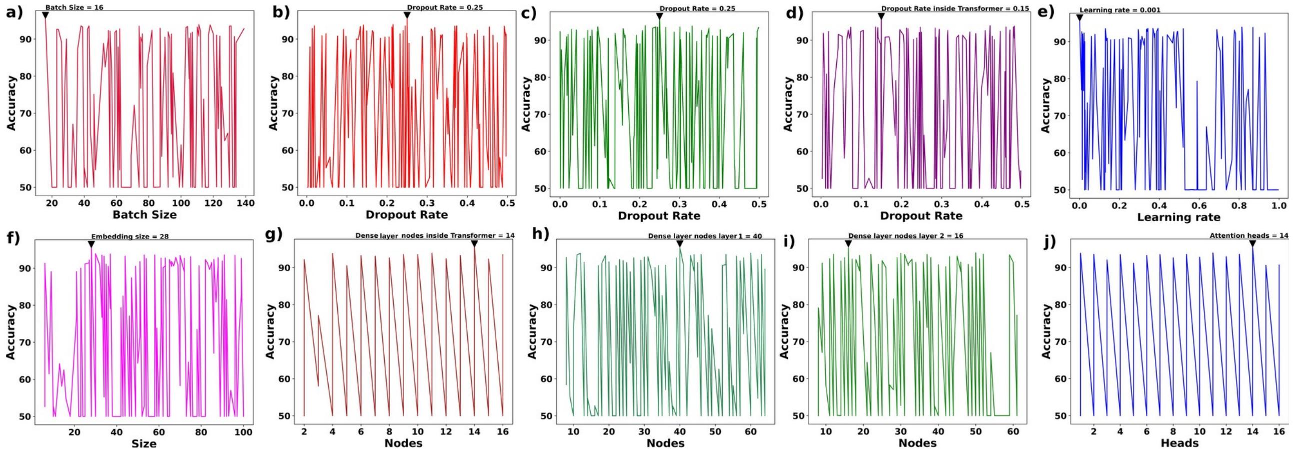


**Supplementary Figure 4: Optimization results for hyperparameters for the transformers model. a)** Batch size optimization, **b)** Dropout rate optimization inside encoder, **c)** First Dropout rate optimization, **d)** Second Dropout rate optimization, **e)** Learning rate, **g)** Embedding size, **h)** Number of units per dense layer inside Transformer, **i)** Number of units per dense layer 1, **j)** Number of units per dense layer 2, and **k)** Number of Attention heads.

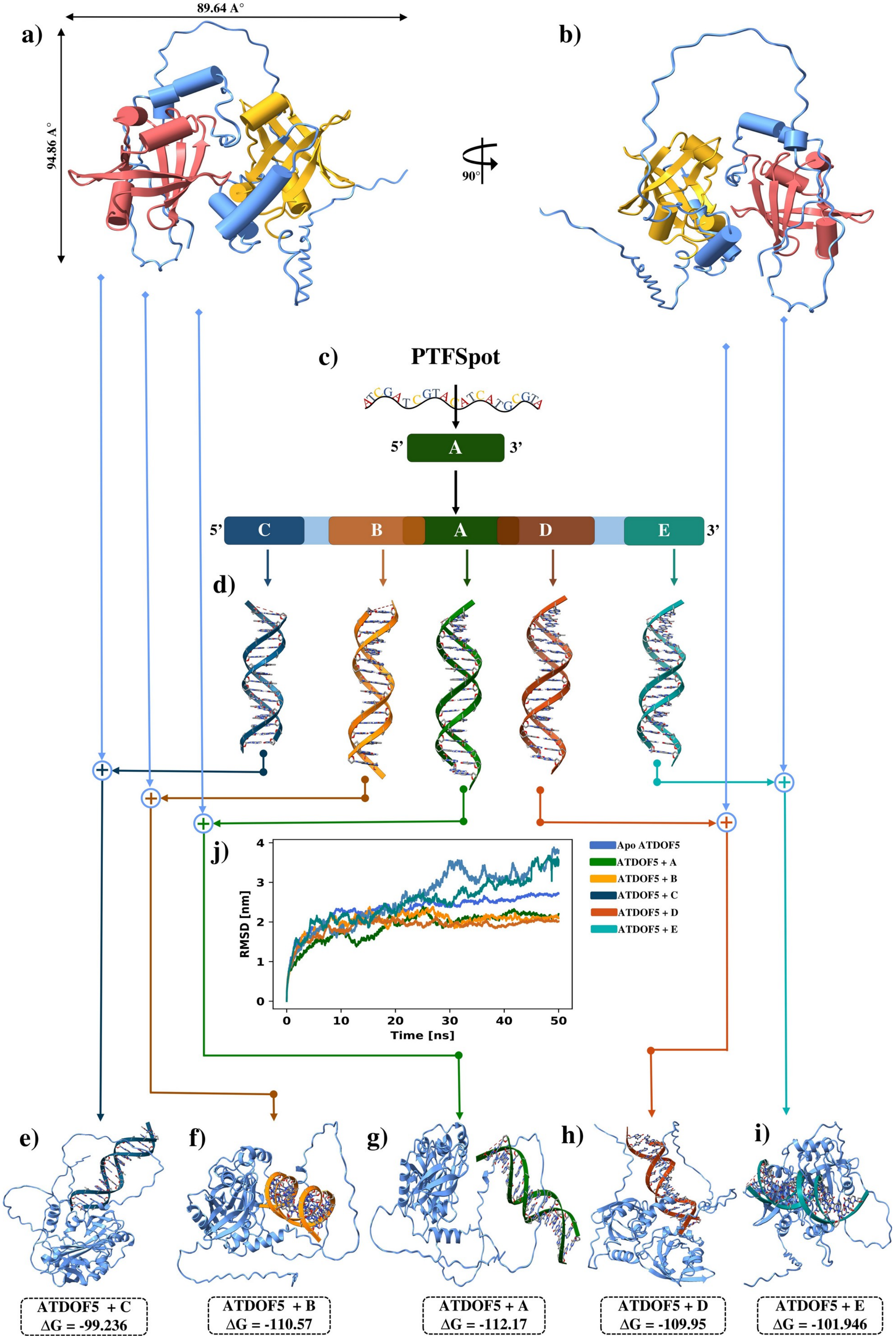
89.64 A°
94.86 A°
a)
b)
90°
c)
PTFSpot
5'
A
3'
5'
C
B
A
D
E
3'
d)
j)
Apo ATDOF5
ATDOF5 + A
ATDOF5 + B
ATDOF5 + C
ATDOF5 + D
ATDOF5 + E
RMSD [nm]
Time [ns]
e)
f)
g)
h)
i)
ATDOF5 + C
ΔG = -99.236
ATDOF5 + B
ΔG = -110.57
ATDOF5 + A
ΔG = -112.17
ATDOF5 + D
ΔG = -109.95
ATDOF5 + E
ΔG = -101.946

**Supplementary Figure 5: Molecular docking and simulation analysis between Transcription Factor (TF) and its binding motif. (a)** 3D structure of ATDOF5 TF (*Arabidopsis thaliana*) with two domains (colored in Pale Red and Golden Red). **(b)** The arrangement of the loops and helices viewed from top after rotation at 90°, **(c-d)** Prime binding motif identification by PTFSpot for the TF ATDOF5, followed by selection of prime binding motif region as well as regions overlapping with it in its flanks and terminal end regions from both sides (A, B, D, C, and E regions). Respective DNA 3D structures were generated for these regions as the central binding one. **(e-i)** The docked pose of each tiling-deletion variant DNA with the TF ATDOF5 and their binding affinity. Out of five docked complexes, the prime motif docked complex showed highest binding affinity than other complexes clearly supporting the importance in TF DNA interactions. **(j)** Molecular dynamics simulation of Apo TF and all five docked complex, the RMSD plot showed that the docked complex of ATDOF5 + region "A" exhibited lowest RMSD value among all, clearly reiterating the above observation that the prime motif region has importance in TF DNA interactions.

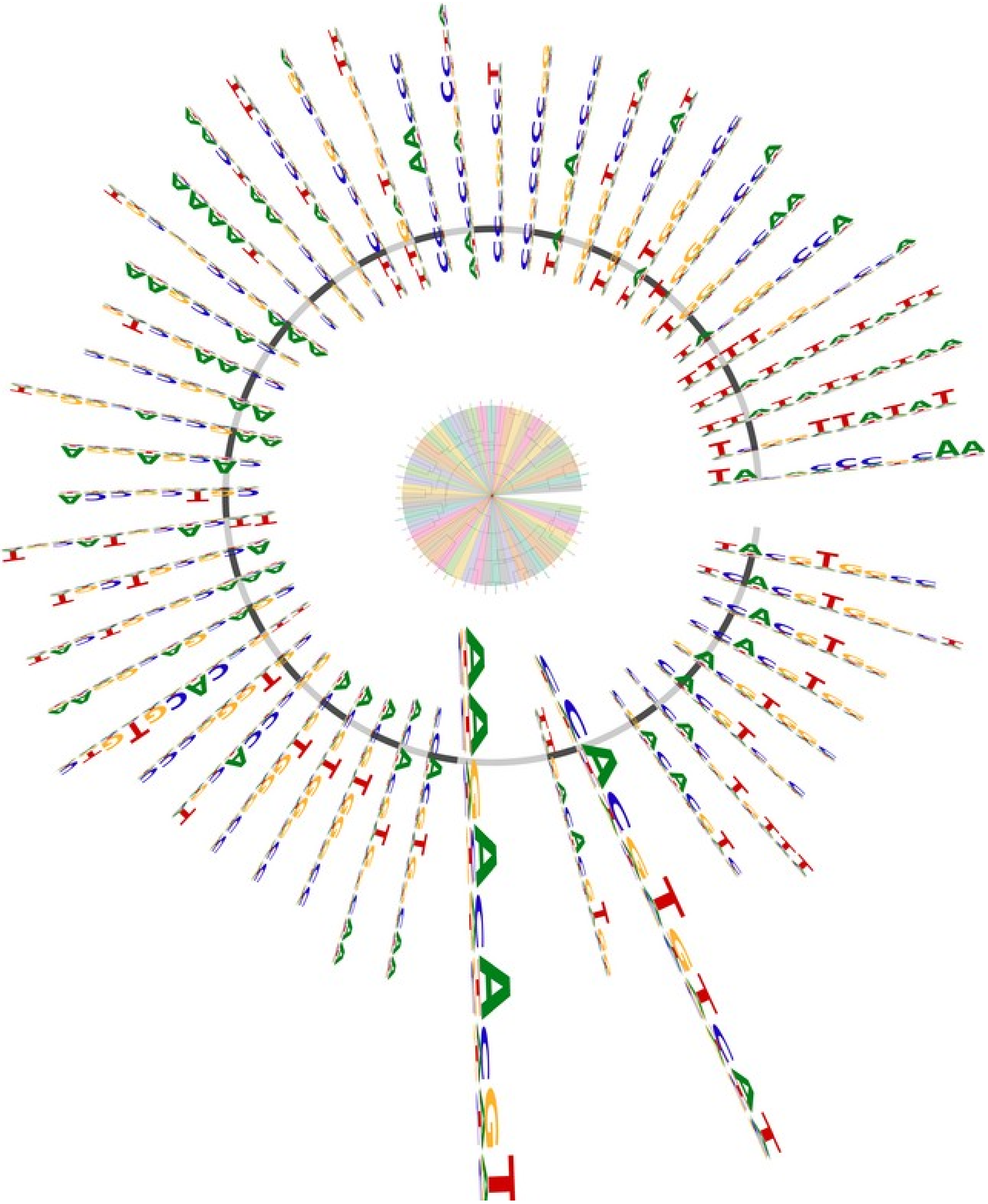

**Supplementary Figure 6: Clustering plot of prime motifs from Dataset "A".** This one covers the prime motifs for 54 transcription factors, and their clustering based on the motifs similarities.

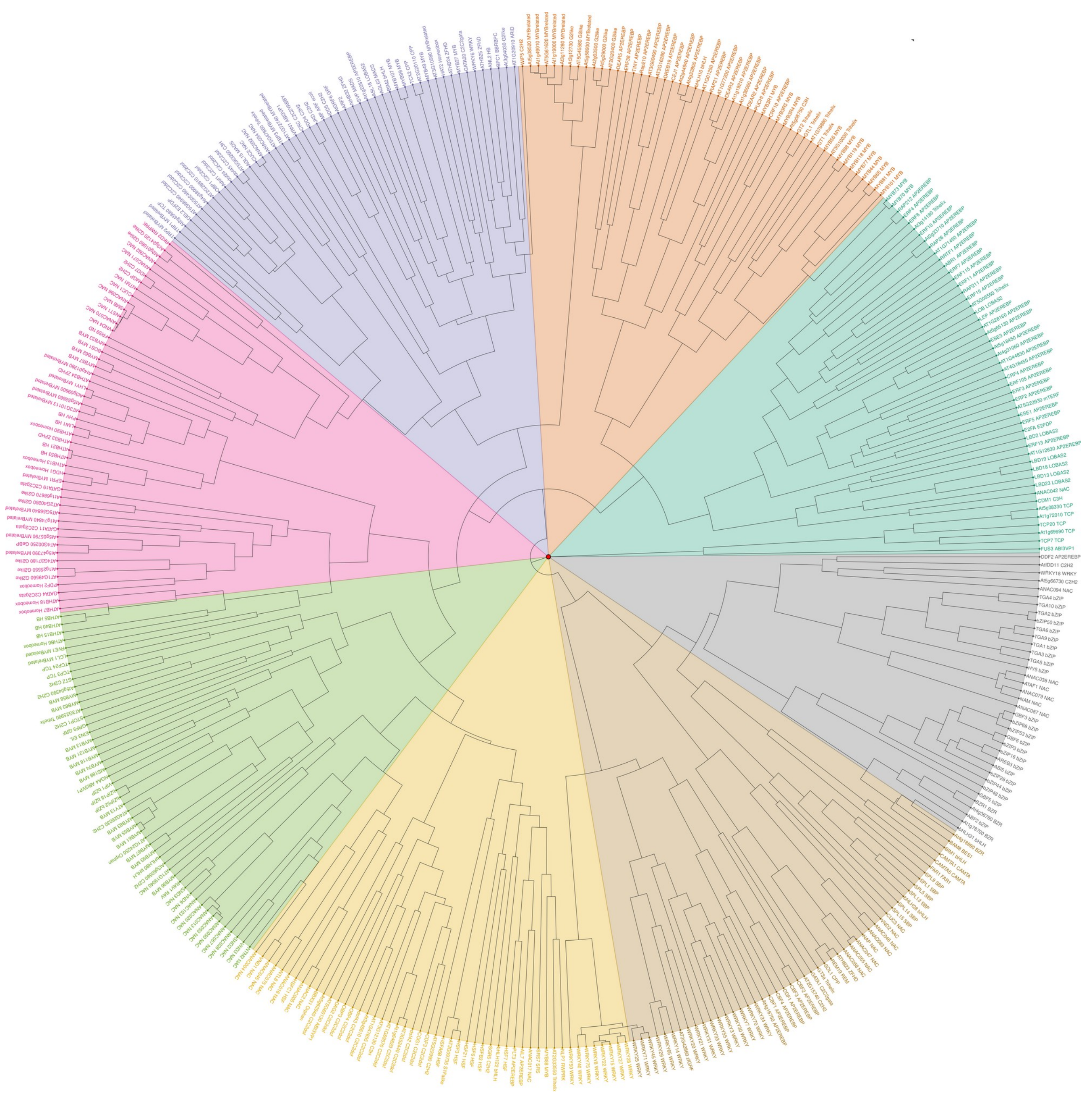

**Supplementary Figure 7: Clustering plot of prime motifs from Dataset "B".** This one covers the prime motifs for 387 transcription factors, and their clustering based on the motifs similarities.

**a)**

<table>
<tr><th>TF</th><th>JASPAR Motif</th><th>Prime Motif</th><th>Alignment</th><th>% of occurence for JASPAR Motif</th><th>% of occurence for Prime Motif</th></tr>
<tr><td>ABF1</td><td></td><td></td><td>ACACGTGGCA<br>||| |||| |||<br>ACACGTGGCA</td><td>72.9</td><td>96.5</td></tr>
<tr><td>ABF3</td><td></td><td></td><td>CACGTGTCTT<br>| ||| ||| |.|<br>CACGTGTCAT</td><td>75.7</td><td>96.1</td></tr>
<tr><td>ABF4</td><td></td><td></td><td>CCACGT<br>||||||<br>CCACGT</td><td>71.5</td><td>96.5</td></tr>
<tr><td>ABI5</td><td></td><td></td><td>TGCCACGTG<br>||. ||||||<br>TGACACGTG</td><td>75.9</td><td>87.2</td></tr>
<tr><td>GBF2</td><td></td><td></td><td>ACGTGTCA<br>| || ||.||<br>ACGTGGCA</td><td>73</td><td>95.2</td></tr>
<tr><td>GBF3</td><td></td><td></td><td>CCACGTG<br>| || ||||<br>CCACGTG</td><td>78</td><td>94.1</td></tr>
<tr><td>KAN1</td><td></td><td></td><td>ATATATT<br>| .| ||||<br>AGATATT</td><td>80</td><td>91.5</td></tr>
<tr><td>PIF1</td><td></td><td></td><td>CCACGTGTCAT<br>| || |||| .|.|<br>CCACGTGACCT</td><td>70.5</td><td>94.7</td></tr>
<tr><td>PIF4</td><td></td><td></td><td>CACGTGT<br>| | ||||.<br>CACGTGG</td><td>90</td><td>94.6</td></tr>
<tr><td>PIF5</td><td></td><td></td><td>CCACGTGA<br>| || ||| ||<br>CCACGTGA</td><td>90</td><td>96.3</td></tr>
<tr><td>SOC1</td><td></td><td></td><td>TTTTGG<br>||||||<br>TTTTGG</td><td>85</td><td>93.6</td></tr>
</table>

**b)**

<table>
<tr><th>TF</th><th>JASPAR Motif</th><th>Prime Motif</th><th>Alignment</th><th>% of occurence for JASPAR Motif</th><th>% of occurence for Prime Motif</th></tr>
<tr><td>TGA4</td><td></td><td></td><td>TGACGTCGT<br>|| |||||.|<br>TGACGTCAT</td><td>94.3</td><td>95.3</td></tr>
<tr><td>GATA19</td><td></td><td></td><td>CCGATCGG<br>|.||||||<br>CAGATCGG</td><td>95</td><td>96</td></tr>
<tr><td>LOB</td><td></td><td></td><td>CTCCGCAG<br>|.|| ||.|<br>CGCCGCCG</td><td>46.48</td><td>92.6</td></tr>
<tr><td>WRKY21</td><td></td><td></td><td>GTCGACG<br>|| |.|||<br>GTCAACG</td><td>10.41</td><td>85.7</td></tr>
<tr><td>LEP</td><td></td><td></td><td>GCCGCGA<br>| ||||.|<br>GCCGCCA</td><td>30.56</td><td>91.2</td></tr>
<tr><td>MYB44</td><td></td><td></td><td>ACGGTTA<br>| ||||.|<br>ACGGTCA</td><td>60.3</td><td>81.5</td></tr>
<tr><td>WRKY29</td><td></td><td></td><td>AACGTCGAC<br>| |.|||.||<br>AAAGTCAAC</td><td>87.35</td><td>85.3</td></tr>
<tr><td>ERF2</td><td></td><td></td><td>CCGACGGC<br>| || .||.|<br>CCGCCGCC</td><td>29.34</td><td>96.1</td></tr>
<tr><td>GATA11</td><td></td><td></td><td>AGATCCG<br>| ||||.|<br>AGATCTG</td><td>98.8</td><td>92.76</td></tr>
<tr><td>TGA9</td><td></td><td></td><td>TGACGTCA<br>|||| ||||<br>TGACGTCA</td><td>83.8</td><td>94.78</td></tr>
</table>

**Supplementary Figure 8: Comparison between experimentally reported motifs and motifs identified in the present study. (a)** Motifs derives from ChIP-seq TF data and (**b)** Motifs derives from DAP-seq TF data. The similarity was determined using TOMTOM package. Several experimentally validated motifs were found similar to the identified prime motifs, with high significance (p-value < 0.01).

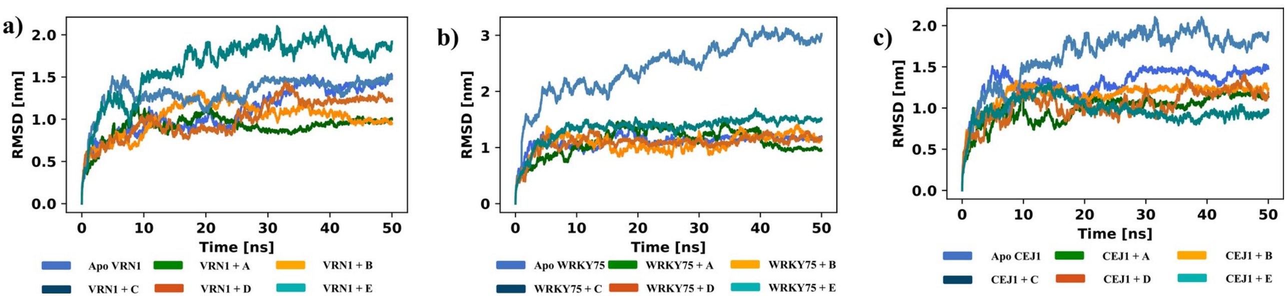


**Supplementary Figure 9: Molecular simulation analysis between the TFs and their respective binding regions for transcription factors a) VRN1, b) WRKY75, and c) CEJ1.**
